# A genetic switch involving EGR1 and EZH2 links aging and development across human tissues

**DOI:** 10.64898/2026.07.29.741427

**Authors:** Neha Khetan, Jiashun Zheng, Changhui Deng, Evan Wu, Hao Li

## Abstract

Aging is often regarded as the last chapter of development. There is growing evidence that aging and development are mechanistically linked. Here we describe a genetic switch involving EZH2 and EGR1 as opposing regulatory nodes that appears to connect aging and development across human tissues. We show that this switch defines two distinct states in fibroblasts: a proliferative (EZH2-high/EGR1-low) state and a non-proliferative and extracellular matrix-expressing (EGR1-high/EZH2-low) state. During replicative aging, cells shift from the proliferative state to the non-proliferative state and targeting either regulatory node—by overexpressing EZH2 or suppressing EGR1—rejuvenates aged fibroblasts. We provide evidence that this switch is involved in the transition from neural progenitor cells to matured neurons during brain development and becomes destabilized during aging, partially reversing the gene expression program established during development. We observed that this same switch becomes blurred and biased towards EGR1-high/EZH2-low state during the aging of muscle stem cells and hematopoietic stem cells. Since genetic switches are widely used in development to establish and reinforce cellular identity, we propose that attenuation of developmental switches may underlie aging across diverse organs and tissues, and partially accounts for the erosion of the epigenetic landscape and the loss of cellular identity.

## Introduction

Development and aging have long been regarded as distinct biological processes and have largely been studied in separate research fields. While developmental biology focus on embryonic development through adulthood, biology of aging has been dominated by the view that the accumulation of molecular and cellular damage is the primary driver. The concept that aging and development are mechanistically linked gained prominence following the discoveries in 1990s that mutations in single genes can substantially extend lifespan, and that these genes were already known to regulate development ^1–4^, suggesting that similar gene regulatory circuits are involved in both development and aging. This has become more apparent from a dynamic systems perspective: the life history of an organism from conception to death may be viewed as a dynamic trajectory governed by the same machinery and gene regulatory networks, and aging is only the later part of that trajectory, a natural continuation of the early part that defines development. Based on this view, developmental models of aging have been proposed that attribute aging to the continuing action or hyper action of developmental programs ^5–11^.

While development establishes the biological programs to build an organism, aging reflects how these programs change and become dysregulated overtime. It is now generally accepted that aging leads to the erosion of the epigenetic landscape established during development to execute and enforce cell fate decisions. Consistently, loss of cell identify during aging was widely observed across different tissues ^12–16^. Such epigenetic drift is considered as a major driving force for aging ^15,17,18^.

Intriguingly, the loss of the order established during development with aging does not appear to be driven by independent, random stochastic processes. Genome-wide DNA methylation pattern seems to change in a predictable way across tissues and organisms, which forms the basis for a universal DNA methylation clock ^19–21^. Transcriptome analysis of aging across tissue/organism revealed groups of functionally related genes consistently decrease or increased with age ^15,22–26^. Systematic drift of gene expression has been observed and correlated with tissue specific age-related diseases ^12^. Interestingly, aging seems to partially reverse the gene expression trajectory of development, where genes up-regulated during development tends to decrease during aging, and vice versa ^27–30^. Together, these observations suggest that age related erosion of epigenetic landscape is not purely stochastic in nature but rather has a collective pattern of change that might be constrained by a gene regulatory network linked to development.

Recent progress in rejuvenation through transcriptional reprogramming has also shed important light on the relation between development and aging. Partial reprogramming with Yamanaka factors (the four transcription factors OCT4, SOX2, KLF4, and MYC, or OSKM) is capable of reversing cellular and tissue-level aging markers and extend lifespan in old mice ^31–34^. Several other single TF perturbations have been found to be able to reverse age associated gene expression programs and rejuvenate cells in vitro and tissues in vivo ^35–40^. In parallel, epigenetic regulators are emerging as promising candidates for cellular reprogramming ^35,41–43^. Interestingly, several of these rejuvenation factors are clearly involved in making developmental decisions. These studies suggest the large-scale epigenetic changes during aging can be reversed by targeting a few key transcription factors or epigenetic regulators, further supporting the notion that age related epigenetic drift is structured, possibly shaped by the gene regulatory network that are involved in making developmental decisions.

These findings suggest that further investigation of the relationship between development and aging may provide valuable insights into the mechanisms of aging and inform the development of rejuvenation interventions. Toward this goal, identifying the genetic circuits that govern key developmental decisions and subsequently drift during aging may be particularly informative, as they enable us to define the underlying dynamics, causal relationships, and functional consequences of the regulators and their downstream targets.

Here we report the identification of a genetic switch involving EZH2 (Enhancer of Zeste Homolog 2), the methyltransferase of PRC2 (Polycomb Repressive Complex 2), and EGR1 (Immediate Early Growth Response 1) that links aging to development across several human tissues. Through systematic perturbation analysis in fibroblasts, we identified a genetic switch with EZH2 and EGR1 functioning as opposing nodes. This switch defines two distinct states, a proliferative state (EZH2+/EGR1-) and a non-proliferative and ECM expressing (EGR1+/EZH2-) state. During fibroblast replicative aging, cells shift from proliferative state to the non-proliferative/ECM expressing state. Over-expressing EZH2 restore the proliferative state and rejuvenate human fibroblasts ^35^.

We provide evidence that the same switch is used in different tissues/organs to regulate transition between proliferative state and non-proliferative but functionally specialized state during development and tissue regeneration. During aging, this switch becomes ambiguous and cells drift away for the state established at young age. For example, during neuro development, as the proliferative neural progenitor cells become mature neurons, cells shift from EZH2+/EGR1- to EGR1+/EZH2-state. During aging, neurons drift away from EGR1+/EZH2-state and populate both. We observed that this same switch becomes blurred and biased towards EGR1+/EZH2-state during the aging of muscle stem cells and hematopoietic stem cells.

Since transition between proliferative vs. non-proliferative and functionally specialized state is a general decision across cell lineages in the context of development and tissue regeneration, we believe EZH2/EGR1 circuit is broadly employed across different tissues. We show that drift of this circuit during aging contributes significantly to the systematic changes of gene expression across tissues linked to diseases ^12^ and can explain the partial reversal of the gene expression programs established during developmental. Since genetic switches are widely used in development to establish and maintain cell identify, we propose that drift of developmental switches might be a major mechanism for the epigenetic drift of aging.

## Results

### I. An EZH2–EGR1 genetic switch defines proliferative and non-proliferative fibroblast states, and its drift underlies replicative aging

In an attempt to systematically identify transcription factor perturbations for rejuvenation, we previously performed a Perturb-seq screening ^35^ (CRISPRa for over-expression and CRISPRi for repression) using a canonical model of human cell replicative aging ^44^. This screen identified a number of single TF perturbations that reverse the gene expression changes during replicative aging, rejuvenating the cells to a younger state. Several TF perturbations (E2F3 and EZH2 over-expression, and STAT3 and ZFX repression) were subsequently validated through molecular phenotyping of aging hallmarks. In particular, EZH2’s rejuvenation effect was validated in vivo in old mouse liver, where overexpression (OE) reverses age associated gene expression changes, reduces fibrosis and steatosis, and improves glucose tolerance ^35^.

EZH2 (Enhancer of Zeste Homolog 2) is the methyl-transferase component of the Polycomb Repressive Complex 2 (PRC2) and plays an important role for gene silencing by methylating H3K27. It is a key developmental regulator and a positive regulator for cell proliferation. It has been observed that during aging, EZH2 level decreases in multiple tissues ^45–49^. We observed decreased expression of EZH2 during fibroblast replicative aging and mouse liver aging, and our perturb-seq analysis shows that EZH2 OE rejuvenates while EZH2 repression accelerates aging in cell-culture and in-vivo in mice, highlighting EZH2 as a key regulator ^35^. Rejuvenation potential of EZH2 OE has also been demonstrated in the aged atrial fibroblasts in mice ^50^ and in the liver fibroblasts of mice and humans ^46^.

To understand better the mechanism of rejuvenation, we examined the top TF perturbations for rejuvenation and accelerated aging for shared themes. While OE of cell cycle regulators (E2F3, FOXM1) or epigenetic silencers (EZH2) rejuvenates, loss of silencing (DNMT1, BRAC1, and EZH2 knock down) gives rise to accelerated aging, indicating the important role of epigenetic silencers for maintaining youthfulness. Intriguingly, repressing TFs with widely different functions (STAT3 for immune response ^51^, ATF4 for integrated stress response ^52,53^, and EGR1 for cell fate decision and stress response ^54,55^) rejuvenates the cells. Of particular interest is EGR1, which ranked first in promoting rejuvenation in the CRISPRi (knockdown) screen and first in accelerating aging in the CRISPRa (overexpression) screen ^35^.

EGR1 (Immediate Early Growth Response 1) is an “immediate early” transcription factor that rapidly responds to external cues to regulate growth, differentiation and stress response ^56^. It regulates different targets in different cellular context. In fibroblast cells, EGR1 mainly regulate genes involved in extra cellular matrix (ECM) remodeling, fibrosis and would healing, including collagens, MMPs and fibronectin ^55,57,58^. Repressing EGR1 leads to rejuvenation with increased proliferation ^59^, consistent with the dichotomy between proliferation vs. ECM deposition modeled previously ^60^. Since we observed increased EGR1 expression (PD32/PD14, log2fc = 0.76, p = 4.4e-6) and decreased EZH2 expression (PD32/PD14, log2fc = -1.04, p = 4.74e-10) during replicative aging ^35^, and perturbing EGR1 and EZH2 in opposite direction generates similar phenotypes, we hypothesized that EZH2 and EGR1 may operate at the opposite ends of a genetic switch to regulate proliferation vs. ECM secretion.

Analysis of the transcriptional response to the top rejuvenating or age accelerating perturbations indeed show the dichotomy of expression of EZH2 and EGR1, as well as markers for cell proliferation and ECM secretion. Rejuvenating perturbations generally lead to increased EZH2 and decreased EGR1, correlated with increased cell proliferation marker such as MKI67 and TOP2A, and decreased expression of ECM genes such as MMP2 and COL1A2. In contrast, TF perturbation that accelerates aging lead to decreased EZH2 and increased EGR1, correlated with decreased cell proliferation markers and increased ECM gene expression **(Figure 1A, Data S1, Figure S1A)**. Similarly, the expression of CDKN2A which encodes the tumor suppressor p16 and p14, showed trend opposite to that of EZH2. CDKN2A is known to be directly repressed by EZH2 ^61^ and has been identified as a longevity gene from genome-wide association studies ^62^.

**Figure 1.**
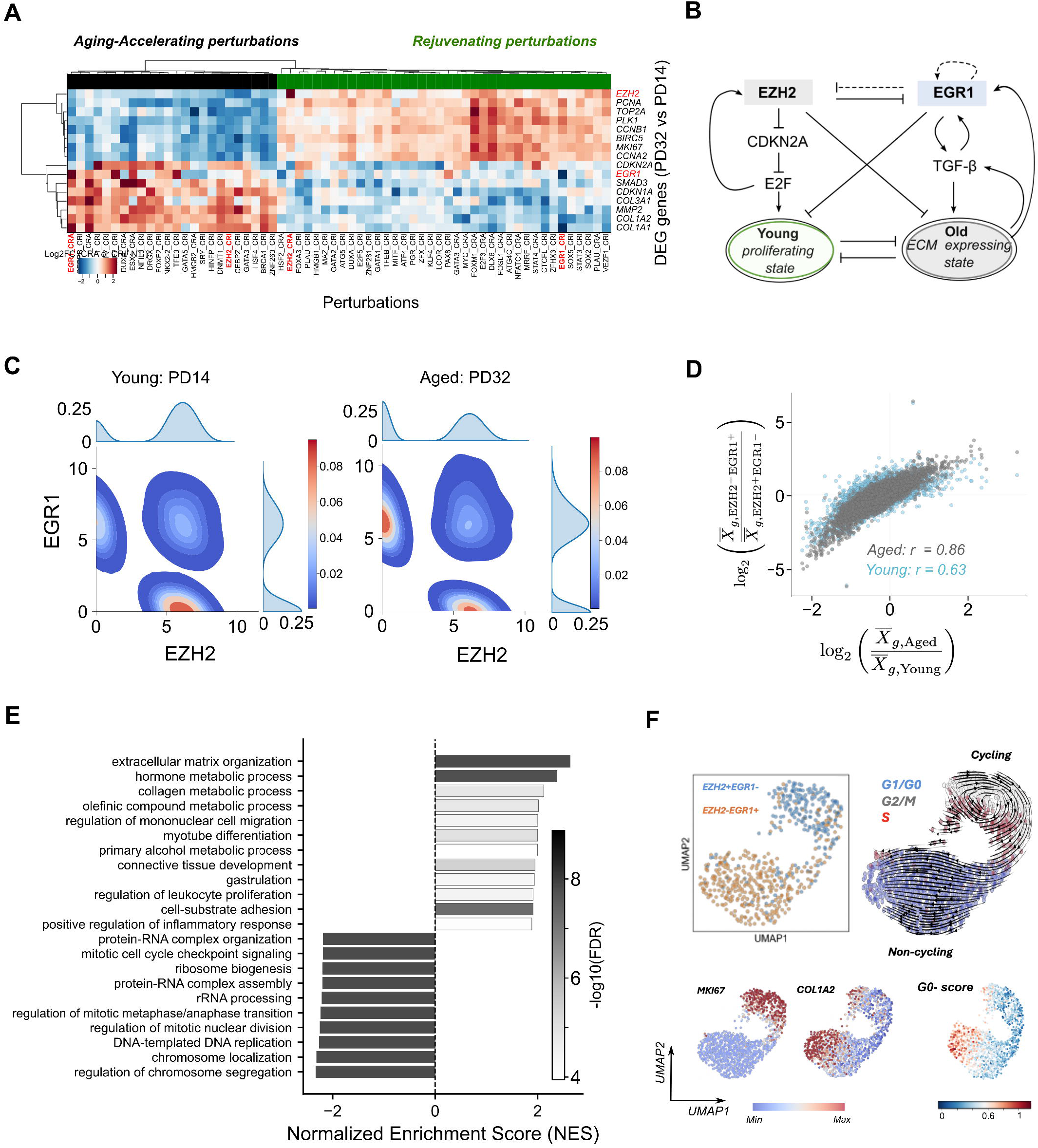
An EZH2–EGR1 genetic switch defines proliferative and non-proliferative fibroblast states, and its drift underlies replicative aging. (A) Dichotomous gene expression programs revealed by perturbation-response analysis. Heatmap showing the expression changes of representative genes, including cell cycle and extracellular matrix (ECM) genes (rows), in response to CRISPRa/i perturbations (columns). Each pixel represents the log2(fold change) in the mean expression of cells carrying the indicated perturbation relative to control cells expressing a non-targeting guide RNA in late-passage fibroblasts (PD32). Perturbations are classified as rejuvenating (*green*) or aging-accelerating (*black*) based on the correlation of their transcriptional responses with differential gene expression observed during replicative aging (PD32 versus PD14). The heatmap was generated by reanalyzing data from *Sengstack* et al. *PNAS 2026*. (B) A simplified gene regulatory circuit illustrating the proposed toggle switch model, with EZH2 and EGR1 functioning as opposing regulatory nodes. This circuit likely represents a core module within a larger gene regulatory network that governs the transition between proliferative and non-proliferative, extracellular matrix (ECM)-expressing states. Solid lines denote interactions supported by direct experimental evidence, whereas dashed lines indicate putative regulatory relationships inferred from studies in other cell types or from bioinformatic analyses. (C) Two-dimensional (2D) density plot of cells in the EZH2–EGR1 expression space in young (PD14) and aged (PD32) fibroblasts. Single cells expressing at least one of the two genes were included. Marginal density plots show the one-dimensional distributions of EZH2 or EGR1 expression across all cells. Gene expression values were normalized and log-transformed, and the 2D density estimates were smoothened (see Methods). (D) Correlation of differential gene expression between the two cell states and the differential expression due to replicative aging. Scatter plots compare differential gene expression between the two subpopulations (EZH2+/EGR1- and EGR1+/EZH2-) within late-passage fibroblasts (PD32, gray) or within early-passage fibroblasts (PD14, blue) with differential gene expression between aged (PD32) and young (PD14) fibroblasts. Each point represents a single gene. (E) Gene set enrichment analysis of Gene Ontology (GO) Biological Process terms associated with genes differentially expressed between the two subpopulations. Bars represent normalized enrichment scores (NES), with color indicating statistical significance. (F) UMAP embedding of fibroblasts showing the EZH2–EGR1 transcriptional states (top left), inferred RNA velocity vectors colored by cell cycle phase (top right), expression of representative marker genes (MKI67 and COL1A2; bottom left and middle), and the calculated G0 cell cycle phase score (bottom right).

Integrating the Perturb-seq data with literature evidence, we propose a simplified gene regulatory circuit in which EZH2 and EGR1 serve as key, opposing regulators with the following basic logic **(Figure 1B)**: 1) EZH2 promote proliferative state while EGR1 promote ECM expressing state, and these two states are mutually inhibitory ^60^; 2) EZH2 positively feedback to itself indirectly through inhibition of cell-cycle inhibitors and tumor suppressor genes (e.g., suppressing CDKN2A ^61^), which in turn suppress positive cell cycle regulators such as E2F family TFs. E2F (particularly E2F1-3) bind to the promoter of EZH2 and positively regulate its expression ^63^; 3) EGR1 positively feedback to itself through TGF-beta pathway activation, ECM secretion and positive regulation by ECM matrix stiffness ^64^, and potentially through self-activation (in-silico motif analysis predicts a self-binding motif ^65^, **Data S2-S3**); 4) EZH2 suppresses EGR1 ^66,67^ and ECM genes directly by silencing their promoters or gene bodies with H3K27me3 markers (EZH2 target genes from ChiP-seq in fibroblasts ^21^ and EZH2 targets in neurons ^68^, **Data S4-S5**), while EGR1 inhibits proliferation by activating cell cycle inhibitors ^54^. EGR1 also regulates microRNAs (e.g., miR-124) that repress EZH2 in neuronal context ^69–71^. Together, these regulatory relationships define a genetic toggle switch with EZH2 and EGR1 acting as key opposing nodes: EZH2 and EGR1 positive feedback to themselves and they mutually inhibit each other, and this circuit controls whether the cell is in a proliferating state or a non-proliferating, ECM expressing state. This circuit is consistent with our systematic perturb-seq data: up-regulation of EZH2, E2F3, and down-regulation of EGR1 all lead to increased cell proliferation and decreased ECM expression (**Figure 1A**), reversing the trends during replicative aging, while down regulation of EZH2 or up-regulation of EGR1 have the opposite effect, leading to accelerated aging.

A genetic toggle switch generally defines two distinct states, and the change at the population level (which could be gradual) due to change of environmental cues can be attributed to the redistribution of cells in the two states ^72–77^. We found that this seems to be the case for the genetic switch described above (**Figure 1B**). This switch defines two distinct states, EZH2+/EGR1- (*S_p_*: proliferating) and the EGR1+/EZH2- (*S_f_*, non-dividing, ECM secreting) states. Young fibroblast (early passage cells) occupied in a clearly defined *S_p_* state and during aging, cells drifted away from the *S_p_* state, with a significant fraction of cells acquired the *S_f_* state (**Figure 1C**).

This shift in the cell population from the *S_p_* to the *S_f_* state largely accounts for the differential gene expression observed between young (early-passage) and old (late-passage) cells, as the transcriptional differences between the *S_p_* and *S_f_* subpopulations within the late-passage cells (or within early passaged cells) largely recapitulate those observed between young and old cells (**Figure 1D, Data S6**). Analysis of the differentially expressed genes between the *S_f_* and the *S_p_* states indicates that up-regulated genes are strongly enriched for ECM organization, wound healing, immune and developmental specification processes, while down-regulated genes are enriched for cell-cycle, DNA repair, mitochondria energy generation, and ribosome biosynthesis (**Figure 1E, Data S7**), supporting the functional switch from a youthful and proliferative state to an old ECM secreting state, consistent with the dichotomy predicted by the model and observed based on candidate markers (**Figure 1A, B**).

We further analyzed the properties of the *S_p_* and *S_f_* states by selecting EZH2+/EGR1- and EGR1+/EZH2-cells and mark them on the UMAP (Uniform Manifold Approximation and Projection) generated for all the cells. These two cell sub-populations clearly separate into two clusters on the UMAP: *S_p_* cells map to the cluster with high MKI67 (proliferation marker) expression but low COL1A2 (collagen) expression, while *S_f_* cells map to the cluster with low MKI67 but high COL1A2 expression (**Figure 1F**). In addition, RNA velocity analysis ^78^ indicates that cells in the *S_p_* state move in closed trajectories, further supporting that they are in active cell cycle and proliferating. For cells in the *S_f_* state, the RNA velocity indicates flow towards cells with high G0 cell cycle phase score, suggesting that they are either in G0 phase or in transition to G0 phase (**Figure 1F**).

Given that *S_f_* and *S_p_* states correspond to “young” and “old” states in the context of fibroblast replicative aging, it is natural to ask whether EZH2 and EGR1 are just passive reporters of the two states or they are the drivers, i.e., shifting of EGR1/EZH2 circuit underlies fibroblast replicative aging? We argue that EZH2/EGR1 circuit is more likely a driver as: 1) EZH2 decreased while EGR1 increased during natural passaging; 2) down-regulation of EZH2 or over-expression of EGR1 (which favor the *S_f_* state, **Figure 1A**) phenocopying aging, causing similar gene expression changes; 3) over-expressing EZH2 or repressing EGR1 (which favor *S_p_* state, **Figure 1A**) rejuvenate fibroblasts; 4) over-expression of EZH2 rejuvenates mouse liver. In addition, the differentially expressed genes between the *S_f_* and *S_p_* states from the late passaged cells are strongly enriched for genes defined as EGR1-responsive from EGR1 over-expression in human dermal fibroblasts and scleroderma patients (227 observed overlapping genes, while expected is 105.67; p:2.84e-34, data from *Bhattacharyya et. al. 2011* ^58^, **Figure S1B**), which are independent studies, supporting the role of EGR1 as a key regulatory node in the gene network that regulates the two states.

### II. The EZH2-EGR1 circuit is involved in regulating the transition from neural progenitor cells to matured neurons during human brain development

The EZH2-EGR1 toggle switch was reconstructed based on perturbation-response analysis of human fibroblast cells. Since EZH2 is the catalytic subunit of PRC2 complex, which is a general epigenetic repressor that have broad function in different tissues, and EGR1 targets different sets of genes in different tissue contexts ^54,56^, we asked whether a similar switch may operate in other cell types to mediate analogous transitions from proliferative cells to non-proliferative, functionally specialized cells.

We turned our attention to the development and aging of neurons in the human frontal cortex, as target prediction based on EZH2 chip-seq data revealed strong enrichment of “transcriptional regulation” and “central nervous system development”, indicating that many of its repression targets are transcriptional regulators for neurogenesis/development (**Data S4 and S5**). Analysis of putative EGR1 target genes, defined by the presence of EGR1-binding motifs surrounding their transcriptional start sites, also revealed strong enrichment for genes involved in neurogenesis and synaptic plasticity (**Data S2 and S3**), suggesting that EZH2 suppresses while EGR1 activates neurogenesis related genes, potentially forming a toggle switch. In addition, EGR1 is known to transcriptionally up-regulate neuron specific microRNAs such as miR-124, which target EZH2 for degradation ^69–71^, and EGR1 is directly repressed by EZH2 ^66,67^ (albeit in different cellular contexts), potentially leading to mutual inhibition that is a key feature of a toggle switch.

The development of the human brain has been characterized using bulk RNA sequencing and, more recently, single-nucleus RNA sequencing (snRNA-seq) ^79–82^. Using the published bulk RNA data ^83^, we found that the dynamics of expression of EZH2 and EGR1 showed opposite trends across different brain regions. EZH2 expression was high during the prenatal period and declined steadily until reaching a plateau at approximately 500 days, whereas EGR1 expression followed the opposite trajectory, increasing from a low prenatal level to reach a similar plateau at approximately the same time (**Figure 2A**). Thereafter, the expression of both genes remained relatively stable throughout adulthood. Concurrently, the cell proliferation marker MKI67 decreased continuously but displayed a two phase dynamics - a rapid decrease phase till ∼120 days followed by a gradual decrease afterward. This dynamics is consistent with the established knowledge in neural development that rapid cell division of neural stem cells mainly happens in the time window from 6 to 17 weeks post conception ^84–86^. Parallel to the increase of EGR1, neuronal maturation and synaptic plasticity markers (such as CAMK2A) increased and reached a plateau around 500 days. Thus, the dynamics of EZH2-EGR1 circuit and the marker genes are consistent with a two phases development: the first phase before 17 weeks post conception with rapid growth and cell division, followed by second phase of slow growth, differentiation, and maturation.

**Figure 2:**
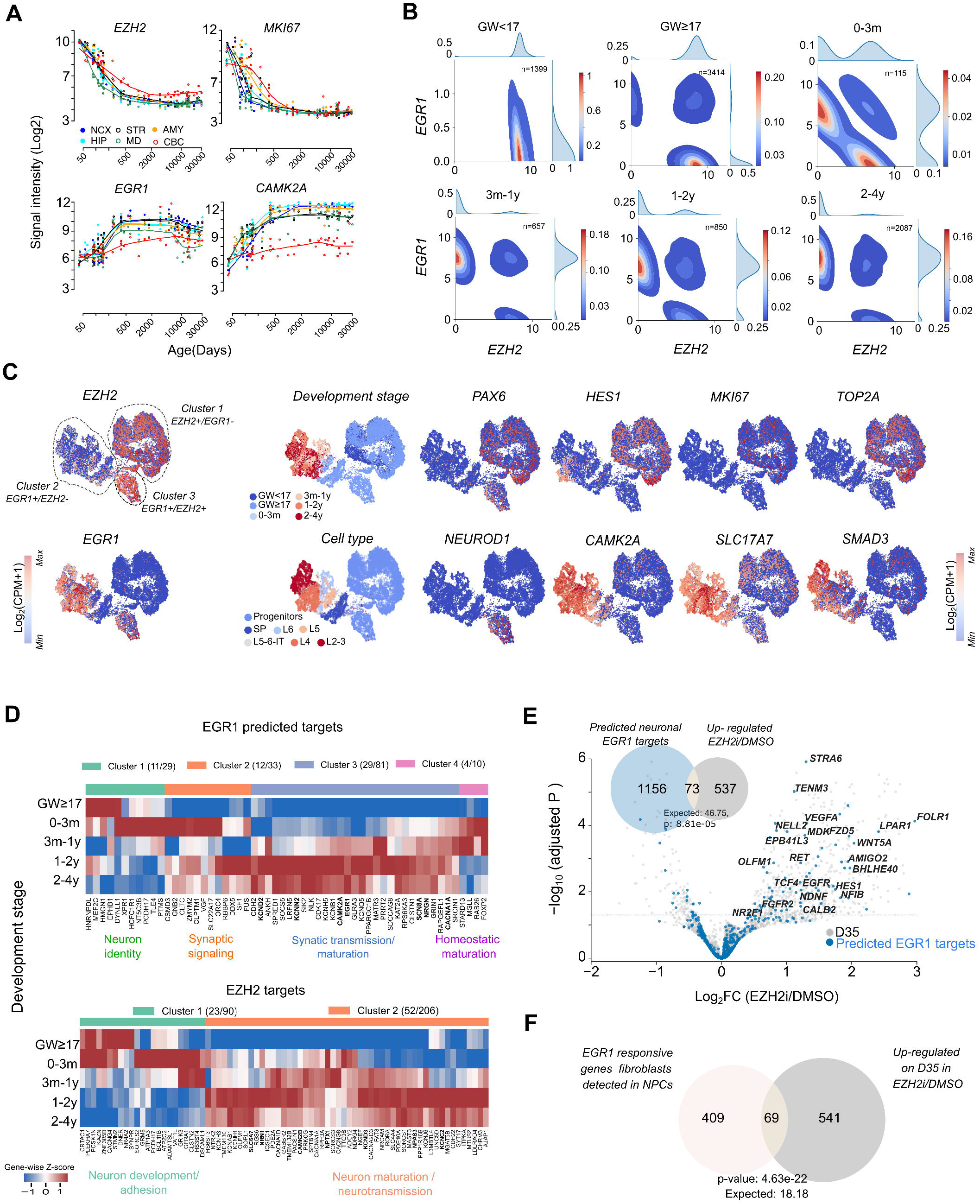
The EZH2-EGR1 circuit is involved in regulating the transition from neural progenitor cells to matured neurons during human brain development. **(A)** Expression of EZH2, EGR1 (*left*), and representative marker genes (MKI67 and CAMK2A) (right) across human brain regions during brain development, measured in microarray assays from brain tissue samples, spanning donors from fetal stage to approximately 80 years of age (age indicated as days post conception). Colors denote brain regions (neocortex: NCX, Cerebellar cortex: CBC, Mediodorsal nucleus of the thalamus: MD, Striatum: STR, Amygdala: AMY, Hippocampus: HIP). Data from the Human Brain Transcriptome (adapted from htttps://hbatlas.org) **(B)** 2D density plots of cells in the EZH2–EGR1 expression space using single nucleus RNA-seq data from human neocortex across different developmental stages, from before gestational week 17 (GW < 17) to 4 years of age. Cells expressing at least one of the two genes were included. Marginal density plots show the one-dimensional distributions of *EZH2* or *EGR1* expression across all cells. Inset indicates total number of nuclei analyzed at each developmental stage. Data reanalyzed from *Velmeshev et. al. Science 2023*. **(C)** UMAP embedding across developmental stages, colored by the expression of marker genes or cell annotations. From left to right: expression of *EZH2* (top) and *EGR1* (bottom), developmental stage (top), and annotated neuronal subtype (bottom). UMAP overlays show the expression of representative marker genes for progenitor/stem cells (PAX6, HES1), proliferating cells (MKI67, TOP2A), differentiating cells (NEUROD1), mature excitatory neurons (CAMK2A, SLC17A7), and ECM/synaptic signaling (SMAD3). **(D)** Heatmaps showing the mean expression of predicted EGR1 target genes (top) and experimentally defined EZH2 target genes (bottom, see Methods) across developmental stages. Genes are grouped by hierarchical clustering (clusters indicated by colored bars), and functional annotations are assigned based on Gene Ontology (GO) enrichment analysis. Representative genes from each functional category are highlighted in bold and top ∼50 variable genes are shown (see, Figure S2 for all genes). Numbers in the parenthesis indicate number of representative genes from and the size of each cluster. *(E)* Volcano plot showing differential gene expression (on day 35) following transient pharmacological inhibition of EZH2 (from day 12 to 20) during in vitro maturation of human neurons. Predicted neuronal EGR1 target genes are highlighted in blue. Inset shows the overlap between genes upregulated following EZH2 inhibition and predicted neuronal *EGR1* targets. Data analyzed from *Ciceri et. al. Nature 2024*. **(F)** Venn diagram showing the overlap between genes upregulated following EGR1 overexpression in human fibroblasts *(Bhattacharyya et. al., PloS One 2011)* and genes significantly upregulated following *EZH2* inhibition in neurons.

Notably, similar expression dynamics were observed across multiple brain regions, suggesting that this phenomenon is not specific to the prefrontal cortex but instead represents a general feature of brain development. It is also interesting to note that cerebral cortex displayed slower kinetics of the marker genes, and this slower kinetics is mirrored in the EGR1/EZH2 dynamics, consistent with the observation that cerebral cortex takes longer to develop than other regions (**Figure 2A**).

Single nucleus RNA-seq data provided higher resolution of different brain regions and cell subpopulations. Analysis of single-nucleus RNA-seq data of the excitatory neurons in the neocortex region during development ^79–82^ indicated that the majority of cells are in two distinct states, consistent with the toggle switch model: an EZH2+/EGR1-state (*S_p_*, proliferative neural progenitor cells) and EGR1+/EZH2-state (*S_f_*, post mitotic, functionally specialized neurons). As development proceeds from prenatal to infancy to toddler, cells shift from *S_p_* to *S_f_*, correlating with the increase of EGR1 and concurrent decrease of EZH2 at the population level, with the distribution stabilized around 1-2 years of age and maintained throughout adulthood. This dynamics is consistent with that from the bulk RNA-seq analysis, with a similar time scale for the establishment of matured neuron population (**Figure 2B**).

The cell states defined by the EZH2–EGR1 circuit showed strong concordance with the developmental stage, spatial location, and functional identity of the cells. Visualizing from a UMAP, we observed 3 major clusters (**Figure 2C**). Cluster 1 corresponds to an EZH2+/EGR1-state and maps predominantly to prenatal progenitor populations located in the subplate (SP), progenitor zones, and cortical layer 6 (L6), which is one of the earliest cortical layers to form. These cells possess proliferation markers (MKI67, TOP2A) and neural progenitor cell markers (PAX6, HES1), have low expression of excitatory neuron marker (SLC17A7), and mostly lack the expression of neural differentiation (NEUROD1), maturation (SMAD3), and synaptic plasticity (CAMK2A) markers. Cluster 2 correspond to EGR1+ /EZH2-cells and map to infant to 4 years and cortical layers L2 to L6. These cells are negative for the proliferation markers (MKI67, TOP2A), neural progenitor markers (PAX6, HES1), and early neural differentiation marker (NEUROD1), but express strong excitatory neuron marker (SLAC17A7), early maturation marker (SMAD3), and synaptic plasticity marker (CAMK2A), indicating they are matured excitatory neurons. Cluster 3 corresponds to EZH2+/EGR1+ cells, and they also map to prenatal progenitor cells. However, they differ from the Clusters 1 cells in that they do not express the proliferative neural progenitor markers (MKI67, TOP2A, HES1), but acquired early neuro differentiation marker (NEUROD1), suggesting that they are in a transition state where NPCs exit the cell cycle and start to differentiate, but has not matured yet. Note that cluster 3 (EGR1+/EZH2+ cells) mainly mapped to progenitor cells > 17 PCW (post-conceptional weeks), while cluster 1 contains both > 17 PCW and <17 PCW progenitor cells. This is consistent with 17 PCW being a key time point beyond which the majority of the NPCs stopped dividing ^85–87^.

To explore the functional relevance of the EZH2/EGR1 switch during brain development, we analyzed the temporal gene expression profiles of predicted EGR1 targets (based on motif analysis, see Methods) and EZH2 targets (based on published datasets on ChIP-seq analysis in fibroblasts ^88^ and H3K27me3 profiling in neurons ^89^, see Methods). EGR1 targets fall into 4 temporal clusters, with cluster 3 being the largest, including ∼80 genes (**Figure S2A and Data S9**). For better visualization, we selected the top ∼50 most variable genes (with a subset of genes in each cluster represented) and plotted their temporal profiles in a heatmap (**Figure 2D).** Consistent with the temporal dynamics of EGR1 at the population level **(Figure 2A)**, The expressions of cluster 3 genes were low in prenatal stage and gradually increased to high level and plateaued in 1 – 2 years. Genes in this cluster are enriched for GO terms “regulation of synaptic plasticity”, “regulation of synapse organization”, “central nervous development”, “regulation of membrane potential”, and “regulation of transcription” (**Data S9**). Cluster 3 genes associated with “regulation of synaptic plasticity” include several well-known neuronal maturation/synaptic plasticity genes such as CAMK2A, which encodes a calcium activated kinase, a widely used marker for excitatory neurons, GRIN1, which encodes an essential core component of NMDA receptor (a ligand gated ion channel responding to glutamate), and NRGN, which encodes Neurogranin that regulates synaptic signaling. The expression of these genes in single cells correlate well with the expression of EGR1, as viewed from the UMAPs (**Figure 2C**). Together, these data suggest as cells switch from EZH2+/EGR1-state (proliferative progenitor) to EGR1+/EZH2-state (differentiating/matured neurons), EGR1 is involved in turning on hundreds of genes important for neuronal maturation, signal transmission between neurons, and synaptic plasticity, arguing that this circuit is actively involved in the key developmental decision.

Analysis of the temporal expression profile of EZH2 target genes added further support to the toggle switch model and functional importance of the switch. EZH2 targets fall into two major temporal clusters, with Cluster 2 being the largest, including **∼**200 genes (**Figure S2B and Data S10)**. The temporal dynamics of the top ∼25% most variable genes were shown in the heatmap **(Figure 2D).** Parallel to the decrease of EZH2 from prenatal to 1-2 years, cluster 2 genes showed an inverted profile, suggesting de-repression as EZH2 goes down. This cluster is enriched for genes related to function of matured neurons **(Data S10),** including SLAC6A7, which encodes a neuron-specific neurotransmitter transporter, and CAMK2B, which encodes a calcium/calmodulin-dependent kinase that regulates synaptic plasticity.

Further evidence supporting that the switch might be causal for the transition from proliferative neural progenitor cells to matured neurons came from studies indicating that EZH2 acts as an epigenetic barrier for neuronal maturation ^89–91^. EZH2 knockdown by the pharmacological inhibitors significantly speeds up the maturation process ^90^. We observed that genes significantly induced with EZH2 inhibition during in vitro maturation significantly overlap with neuronal genes predicted to be EGR1 targets **(Figure 2E).** The overlapping genes are enriched for neuronal-fate determination, axon formation and neuron migration (**Data S11**), suggesting the down regulation of EZH2 during neuron maturation drives the de-repression of functional neuronal genes necessary for the establishment of matured neuronal state. Furthermore, genes induced by EZH2 inhibition strongly overlap with genes induced by EGR1 over-expression in fibroblasts ^58^ (**Figure 2F, Data S12**), arguing that EZH2 and EGR1 function in opposite directions in neurons, and that there is certain conservations of EGR1 targets across tissues, even though the EGR1-EZH2 circuit might be modified by lineage specific regulators.

### III. Drift away from the developmentally established EGR1+/EZH2-state contributes to neuronal aging and partial reversal of the developmental gene expression programs

Given the role of the EZH2–EGR1 circuit in brain development and our finding that drift of this circuit accompanies replicative aging in fibroblasts, we asked whether a similar drift might also contribute to neuronal aging. In a recent genome-wide study characterizing transcriptome and methylome changes at single cell level due to brain aging, EGR1 binding sites were found to be enriched in identified differentially hyper-methylated regions in aged neurons, compared to neurons from younger individuals ^92^. This was attributed to reduced TET1 demethylase recruitment to the specific target genes due to decreased EGR1 expression ^92–95^. The transcript level of EGR1 was also observed to be significantly down regulated in different neuronal types across dorsal lateral pre-frontal cortex (DLPFC). Several other studies also point to the importance of EGR1 for neuronal aging ^92,96,97^.

We have analyzed the potential drift of the EZH2-EGR1 circuit and its functional consequences using several published single cell brain aging datasets, including data published by Chien et. al. ^92^ and a more recently published dataset from a larger cohort by Catching et al. ^98^. We found a consistent pattern of drift: in young subjects, neurons populated the EGR1+/EZH2-state, which is a functional “young” state established after neuronal maturation. During aging, neurons drifted away from the young state and a significant fraction of the neurons acquired the “old”, EZH2+/EGR1-state (**Figure 3A**). Note that the direction of the drift is opposite to that for fibroblasts, which drifts away from the EZH2+/EGR1- (the young and proliferative state) state during aging (**Figure 1C**).

**Figure 3:**
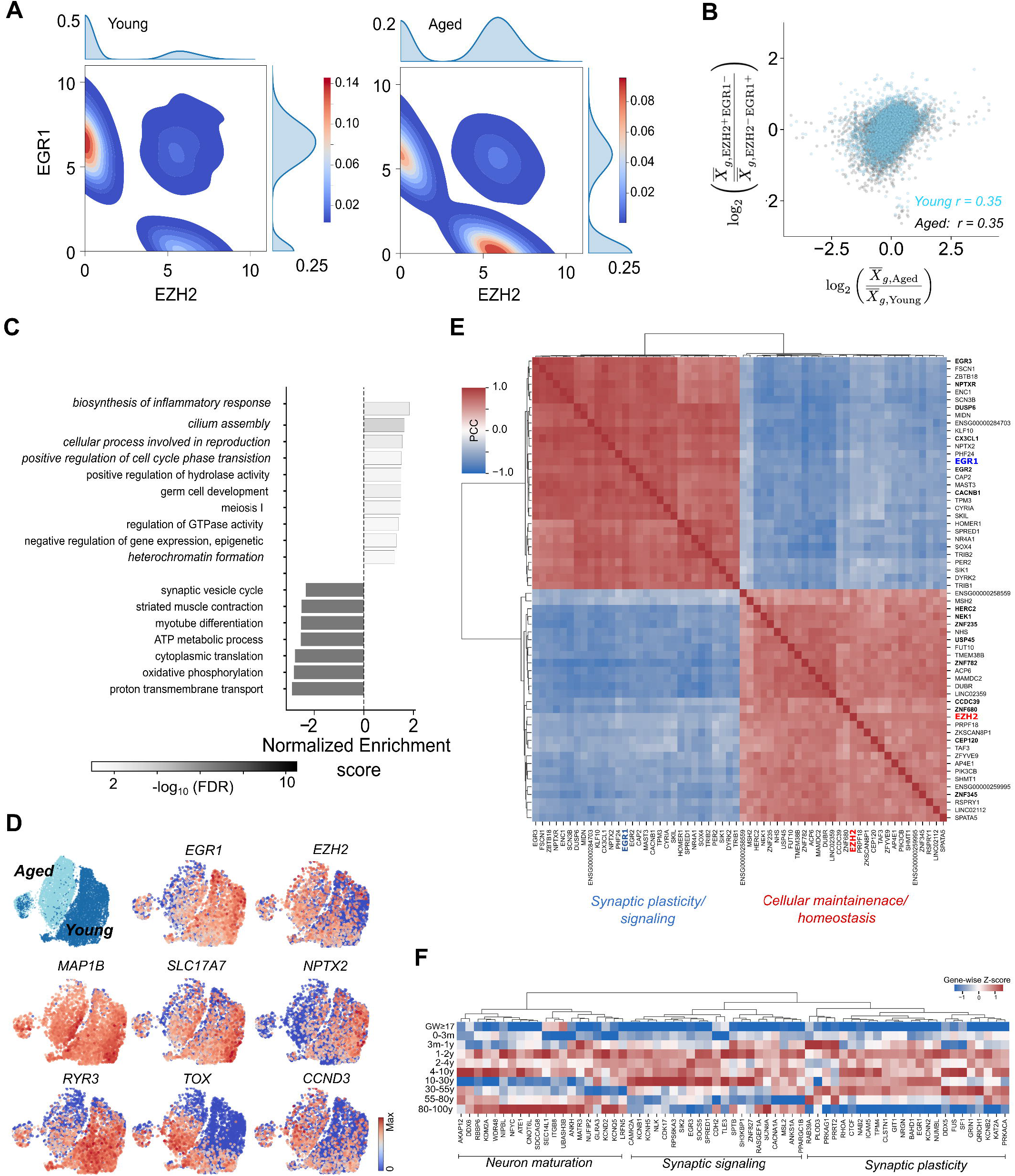
Drift away from the developmentally established EGR1+/EZH2-state contributes to neuronal aging and partial reversal of developmental gene expression programs. **(A)** 2D density plots of cells in EZH2 and EGR1 expression space in upper-layer excitatory neurons (L2–L3) from young (10–30 years; 14 donors; 2,961 cells) and aged (>80 years; 17 donors; 1,210 cells) female donors. Cells expressing at least one of the two genes were counted. For the marginal distribution of EZH2 or EGR1, all cells were counted. Data reanalyzed from *Catching et al. Cell Rep.* 2026. **(B)** Correlation between age-associated differential gene expression and differential expression between the two cell subpopulations (EZH2+/EGR1- and EGR1+/EZH2-). Scatter plots compare age-associated differential expression with differential expression between the two cell subpopulations within aged (gray) and young (blue) excitatory neurons. Each point represents a single gene. **(C)** Gene set enrichment analysis showing significantly enriched Gene Ontology biological processes associated with genes differentially expressed between the EZH2+/EGR1- and EGR1+/EZH2-subpopulations in the aged samples. Bars represent normalized enrichment scores (NES), with color encoding significance. **(D)** UMAP embedding showing cells from young and aged donors together with the expression of EGR1, EZH2, and some representative genes associated with neuro-transmission and synaptic plasticity (SLC17A7, MAP1B, NPTX2, RYR3), cell cycle (CCND3), and T cell/neuronal development (TOX). **(E)** Heatmap of pairwise Pearson correlations among the top 20 genes positively correlated with EGR1 and the top 20 genes positively correlated with EZH2. Correlations were calculated using pseudo-bulk transcriptomes across female donors aged 10–100 years. Hierarchical clustering identified two major transcriptional modules, corresponding to synaptic plasticity and stress remodeling/neuronal homeostasis. **(F)** Heatmap showing the mean expression of predicted neuronal EGR1 target genes across developmental and aging stages. Genes are grouped by hierarchical clustering, with functional categories assigned based on Gene Ontology enrichment analysis.

The shifting of cell population from EGR1+/EZH2- to EZH2+/EGR1-state contributes significantly to the differential gene expression between old and young neurons. Differential expression between the two sub-populations of the old cells significantly correlates with the differential expression between the old and the young cell populations. Similar correlation is observed when the two subpopulations were taken from the young cells (**Figure 3B**). This suggests that a substantial fraction of the seemingly continuous age-associated changes in gene expression arises from a redistribution of cells between these two states.

Functional analysis of genes differentially expressed between the “old” EZH2+/EGR1- and the “young” EGR1+/EZH2-states indicates that genes normally repressed in the young state are up-regulated in the old state, including those related to inflammation and cell cycle, while genes having important functional role in neurons are down-regulated, including those related to synaptic plasticity, mitochondrial function/ATP synthesis, and protein synthesis (**Figure 3C, Data S13**). The up and down regulated genes are also consistent with what is known about the targets of EZH2 and EGR1 - the former represses tumor suppressor genes such as CDKN2A and CDKN1A and thus positively regulate cell cycle, while the latter positively regulates synaptic plasticity and other functional neuronal genes.

To further characterize the two states defined by the EZH2-EGR1 circuit, we visualized the cells by UMAPs generated from the full transcriptome data. The EGR1+/EZH2- and the EZH+/EZH2-cells are separated on the UMAPs, with strong overlap with the young and old samples respectively (**Figure 3D**). Cells in both states have the canonical neuronal markers such as MAP1B and SLC17A7. However, other important functional genes such as NPTX2 (regulating excitatory synapse) and RYR3 (intra-cellular calcium release channel) are highly expressed in the EGR1+/EZH2-state but significantly decreased in the EZH2+/EGR1-state. In contrast, other genes not needed in mature neurons are aberrantly expressed in the EZH2+/EGR1-state, such as TOX for T-cell development and CCND3 for regulation of cell cycle. Interestingly, cells in both states do not express cell proliferation markers (such as MKI67, TOP2A; **Data S14**), suggesting that cell proliferation is under tight control even though old neurons express some of the cell cycle genes.

A global analysis of correlation between genes across individuals based on pseudo-bulk data also suggests that EGR1 and EZH2 represent two groups of covarying genes that function antagonistically, possibly forming a larger gene regulatory network that defines the two states. Genes positively correlated with EZH2 and those positively correlated with EGR1 form two “modules” that are strongly negatively correlated with each other, displaying a dichotomic pattern of expression (**Figure 3E**). Consistent with functional analysis of the two states, the EGR1 modules are enriched for functional neuronal genes such as genes for synaptic plasticity (NPTX2, NPTXR, HOMER1) and ion channels (CACNB1 SCN3B), while the EZH2 module are enriched for DNA repair (USP45, HERC2, MSH2, NEK1, YLPM1), centrosome/cilia biology (CCDC39, CEP120), and zinc finger transcription factors, mostly not functionally characterized (ZNF345, ZNF235, ZNF782, ZNF680).

The drift of the circuit away from the EGR1+/EZH2-state during neuronal aging is opposite to the direction of the change during development/neuron maturation, where the circuit switches from EZH2+/EGR1- (proliferative neural progenitor) state to EGR1+/EZH2- (matured neuron) states. This is consistent with the notion that aging to a certain degree reverses the developmental program and erodes the epigenetic landscape established during development ^27–30^.The functional consequences of this drift are reflected in the significant decline in the expression of predicted EGR1 target genes during aging; many of them are canonical functional neuronal genes involved in synaptic plasticity and learning and memory (E.g., CAMK2A, KCNB1, NRGN, CLSTN1, GRIN1, SCN8A) (**Figure 3F**). The expression of these genes were established as neural progenitor cells developed into matured neurons but got lost in old age as the cells shift away from the EGR1+/EZH2-state established at the young age. Thus the drift of the circuit with aging may lead to decreased expression of functional neuronal genes that drive the decline of memory and executive function.

### IV. EZH2-EGR1 switch becomes blurred/biased towards the EGR1+/EZH2-state with hematopoietic and muscle stem cells aging

Having observed age associated drift of EZH2-EGR1 switch in both proliferative fibroblasts and post-mitotic neurons, we next asked whether this circuit exhibits similar drift in tissue-specific stem cells, which dynamically transition between quiescent, proliferative, and post-mitotic differentiated states throughout life. EZH2 is generally highly expressed in many types of adult stem cells and progenitor cells across tissues ^49,90,99–101^, and EGR1 is known to be an important regulator of hematopoiesis and myeloid lineage differentiation ^102–104^ **(DataS2 and Data S3)**. We thus explored whether drift of this circuit is implicated in HSC aging.

HSCs transition from quiescent, long-term HSCs (LT-HSC) (GATA2+, FOXO3+) to short term, proliferative HSCs (ST-HSC) (MKI67+), to multi-potent progenitor (MPP) cells which then differentiate into Common Lymphoid Progenitor (CLP) (IKZF1+) lineage and common myeloid progenitor (CMP) (CLU+) lineage to produce lymphoid cells and myeloid cells (**Figure 4A**). Analyzing single-cell transcriptome data of donors from the published atlas (*Li. et. al.* ^105^) across lifespan (pre-neonates:10-23 gestational weeks, pediatric:2-12 years, adult:17-53 years and aged: 62-77 years), we observed a systematic shift of cells in the EZH2-EGR1 expression plane. HSCs in fetal samples populated the EGR1+/EZH2-state, suggesting there are more quiescent than proliferative cells (**Figure 4B**). In the cord blood, more cells switched to the EZH2+/EGR1-states, indicating transition to more proliferative state to rapidly expand the HSC pool. As development proceeds to pediatric and adult stages, a more balanced distribution between the two states is reached. Interestingly, aging disrupts this balance, shifting the distribution toward the EGR1+/EZH2− state, suggesting that HSCs increasingly adopt a more quiescent state (**Figure 4B**).

**Figure 4:**
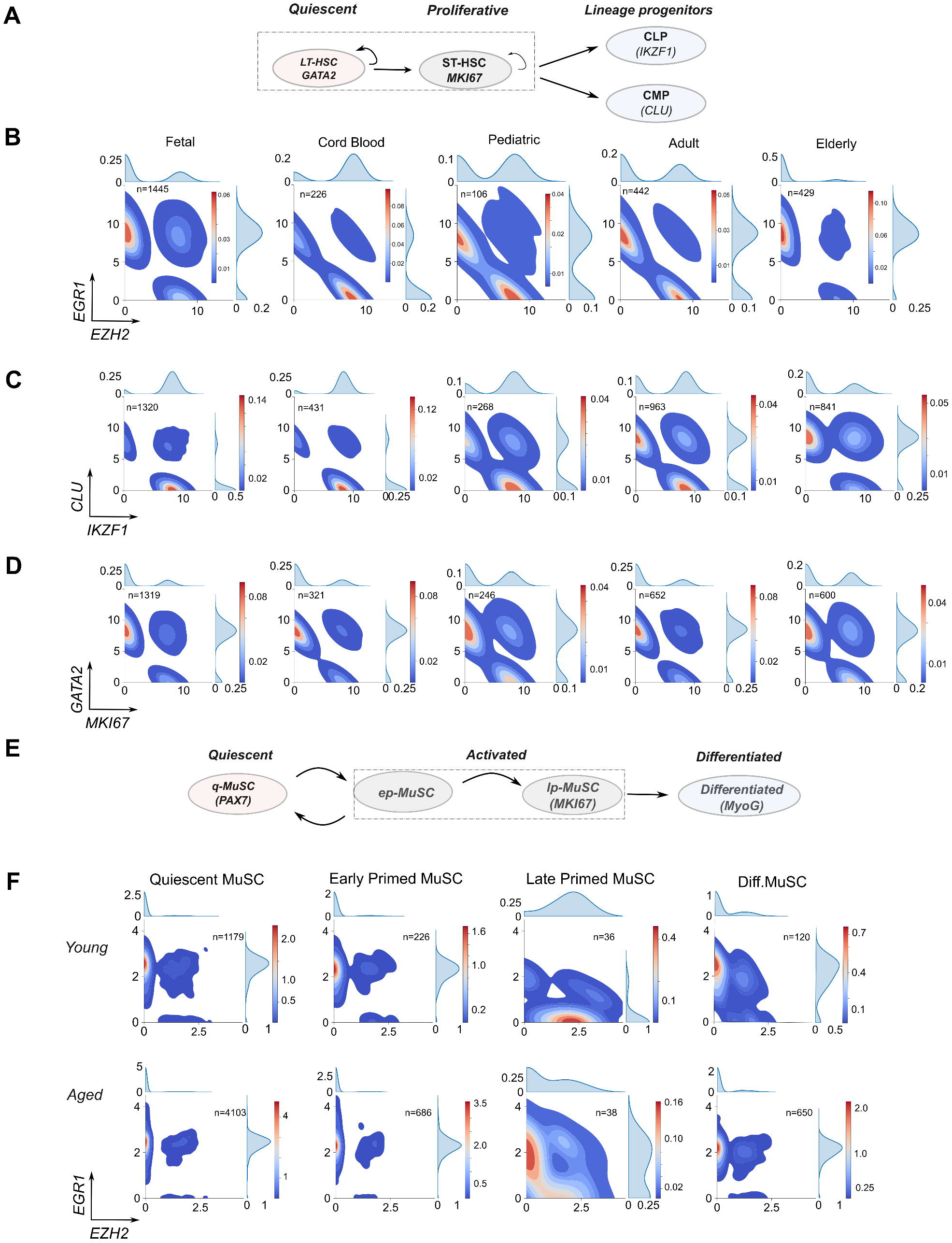
EZH2-EGR1 switch becomes blurred/biased towards the EGR1+/EZH2-state with hematopoietic and muscle stem cells aging. **(A)** Schematic illustrating hematopoietic stem cell (HSC) state transitions from long-term quiescent HSCs (LT-HSCs) to proliferative short-term HSCs (ST-HSCs), and subsequently to lineage-committed multipotent progenitors, including common lymphoid progenitors (CLPs) and common myeloid progenitors (CMPs). **(B)** 2D density plots of single HSCs in the EZH2–EGR1 expression space across developmental and aging stages (fetal, cord blood, pediatric, adult, and elderly). Cells expressing at least one of the two genes were included in the density plots (cell numbers are indicated in the inset). Histograms show the marginal distributions of EZH2 or EGR1 expression across all cells. Data from *Li et al. Nat Methods 2025*. **(C-D)** 2D density plots showing the expression of lineage and cell-state markers in single-HSCs across the developmental and ageing stages. **(C)** Lineage specific markers: lymphoid (IKZF1) and myeloid (CLU). **(D)** Cell state markers: quiescent (GATA2) and proliferative (MKI67). Cells expressing at least one of the two genes were included (Inset indicates the total number of cells in each stage). Marginal histograms show the expression distributions of each marker across all cells. **(E)** Schematic depicting muscle stem cell (MuSC) state transitions from quiescent (qMuSC), to activated early primed (ep-MuSC) and late primed (proliferative, lp-MuSC), to differentiated myogenic cells (Diff. MuSC). **(F)** 2D density plots of MuSCs in the EGR1-EZH2 expression space across MuSC states in young (7 donors, 15-45 years of age, top panel) and aged (10 donors, 77-99 years of age, bottom panel) MuSCs. Cells expressing at least one of the two genes were included (Inset indicates the total number of cells for each state). Marginal histograms show the expression distributions of each gene across all cells. Data from *Lai et. al., Nature 2024*.

Consistent with the above interpretation, we observed similar shift of distribution in the GATA2 and MKI67 expression plane, the former marks the quiescent state of LT-HSC, while the latter is a marker of cell proliferation. Cells occupy more GATA2+/MKI67-state in the fetal stage, reach a more balanced distribution in the Pediatric stage, and become biased towards GATA2+/MKI67-state with age (**Figure 4C**). Viewed from the plane defined by CLU and IKZF1, lineage markers for CMP and CLP respectively, cells shift from a balanced distribution in young samples (Pediatric and Adult) and become biased towards CMP+/CLP-state with age (**Figure 4D**), consistent with the observation that myeloid lineage bias is a major hallmark of HSC aging ^105^ ^106^.

Based on the above observation, we hypothesize that the drift of the EZH2-EGR1 circuit with age towards EGR1+/EZH2-state might be an upstream cause for decreased self-renewal and increased myeloid lineage bias. This hypothesis is consistent with previous studies that implicated EGR1 and EZH2 in regulating hematopoiesis and lineage differentiation. EZH2 is essential for long term HSC renewal and lymphopoiesis ^107^, and its loss leads to de-repression of developmental genes including EGR1 ^67^, promoting myeloid bias and associated malignancies ^107^. Furthermore, EGR1 promotes HSC quiescence and bone marrow retention ^103^, and high EGR1 observed to correlate with age associated increase myeloid-biased differentiation, and inflammatory activation ^102,104,108^. In addition, aged HSCs that exit quiescence in response to stress exhibit impaired cell-cycle progression, together with expansion of phenotypic HSC pool but reduced functional self-renewal ^102,109–112^.

To see if the involvement of the EZH2-EGR1 circuit in tissue specific stem cells is more general, we examined whether age-associated drift of the EZH2–EGR1 circuit extends to skeletal muscle satellite cells (MuSCs), the stem cells responsible for skeletal muscle repair and regeneration. Unlike HSCs, MuSCs remain predominantly quiescent under homeostatic conditions and are activated transiently following injury. Upon activation, quiescent MuSCs (PAX7+) progress through an early-primed state enriched for stress-response genes (EGR1+), followed by a late-primed proliferative state (MKI67+), with a subpopulation subsequently undergoing myogenic differentiation (MYOG+) (**Figure 4E**).

Analysis of single-cell/single-nucleus MuSC data from young (15-46 years) and aged donors (74-99 years) ^113^ revealed a similar age-associated redistribution of cells in the EZH2–EGR1 plane (**Figure 4F**). The transition from the EGR1+/EZH2-state in early-primed cells to the EZH2+/EGR1− state in late-primed cells was cleaner in MuSCs from young donors than those from aged donors. Whereas young MuSCs underwent a sharp transition between the two states, aged MuSCs exhibited a predominant population of cells “stuck” in the EGR1+/EZH2-state. Similar pattern was observed in the plane defined by MKI67 and MYOG, two markers representing proliferation and differentiation, where aged MuSC acquired less proliferation marker and retains more differentiation marker at the early and late primed stages (**Figure S3).** Interestingly, MYOG is a known downstream target of EGR1 ^114^, suggesting that aging induced stress perhaps drives higher EGR1 expression that entrench the quiescent non-proliferative state.

## Discussion

There is growing evidence that aging and development are mechanistically linked. Previous theories have proposed that aging represents a continuation or dysregulation of developmental programs ^5–11,23^. Here, we extend this concept by proposing that aging is driven by the progressive loss of fidelity of developmental genetic switches, causing discrete cell states established during development to become increasingly blurred.

We describe a concrete genetic switch involving EZH2 and EGR1 functioning as opposing regulatory nodes that appears to link aging with development. We show that this switch defines two distinct states in fibroblasts and targeting either regulatory nodes can lead to rejuvenation of replicatively aged cells. We provide evidence this switch is involved in the transition from neuro progenitor cells to matured neurons during brain development, and that the same switch becomes ambiguous during aging, partially reversing the gene expression program established during development. We observed that a similar switch becomes attenuated during the aging of HSCs and MuSCs. As genetic switches are widely used in development to establish and reinforce cell identity, we propose that attenuation of these switches may underlie aging across different organ/tissues, accounting for the erosion of epigenetic landscape and loss of cell identify.

A key developmental decision to make across cell lineages is to transition from proliferative progenitor cells to non-dividing and functionally specialized cells. We hypothesize that the EZH2-EGR1 circuit is generally involved in such decision making. EZH2 is the key enzymatic component of PRC2 which is known to play an important role during development and cell fate decision. EZH2 targets many transcription factors and is positively correlated with cell proliferation in fibroblasts (**Figure S1A**, EZH2 and MKI67 are positively correlated across all perturbations in **Figure 1A**, r=0.68, p =1.45e-9), in neural progenitor cells, and in HSC and MSC during regeneration. EGR1 generally acts as a repressor of cell proliferation, possibly indirectly through the induction of tumor suppressors (such p21, p16), TGF-beta, ECM etc. EGR1 targets are also strongly enriched for TFs, and it positively regulates a broad spectrum of functional genes, including genes related to synaptic plasticity, myeloid differentiation, and muscle structure development (**Data S2 and S3**), indicating it is a master regulator. A plausible model is that EZH2-EGR1 switch may operate as part of nested switches that define developmental trajectories (**Figure 5**). In the context of different cell lineages, EZH2-EGR1 switch is employed to make decision to stay proliferative, or to transition to post-mitotic and functionally specialized state for that lineage; the lineage specificity may be achieved through the EZH2-EGR1 circuit molded by the lineage specific regulators. In the context of tissue regeneration (such as in MuSC or HSC), transition between the two states can be triggered by external cues such as injury and stress response. It will be important to investigate the role of EZH2-EGR1 circuit during development and aging in other organs/tissues to see how general this model is.

**Figure 5.**
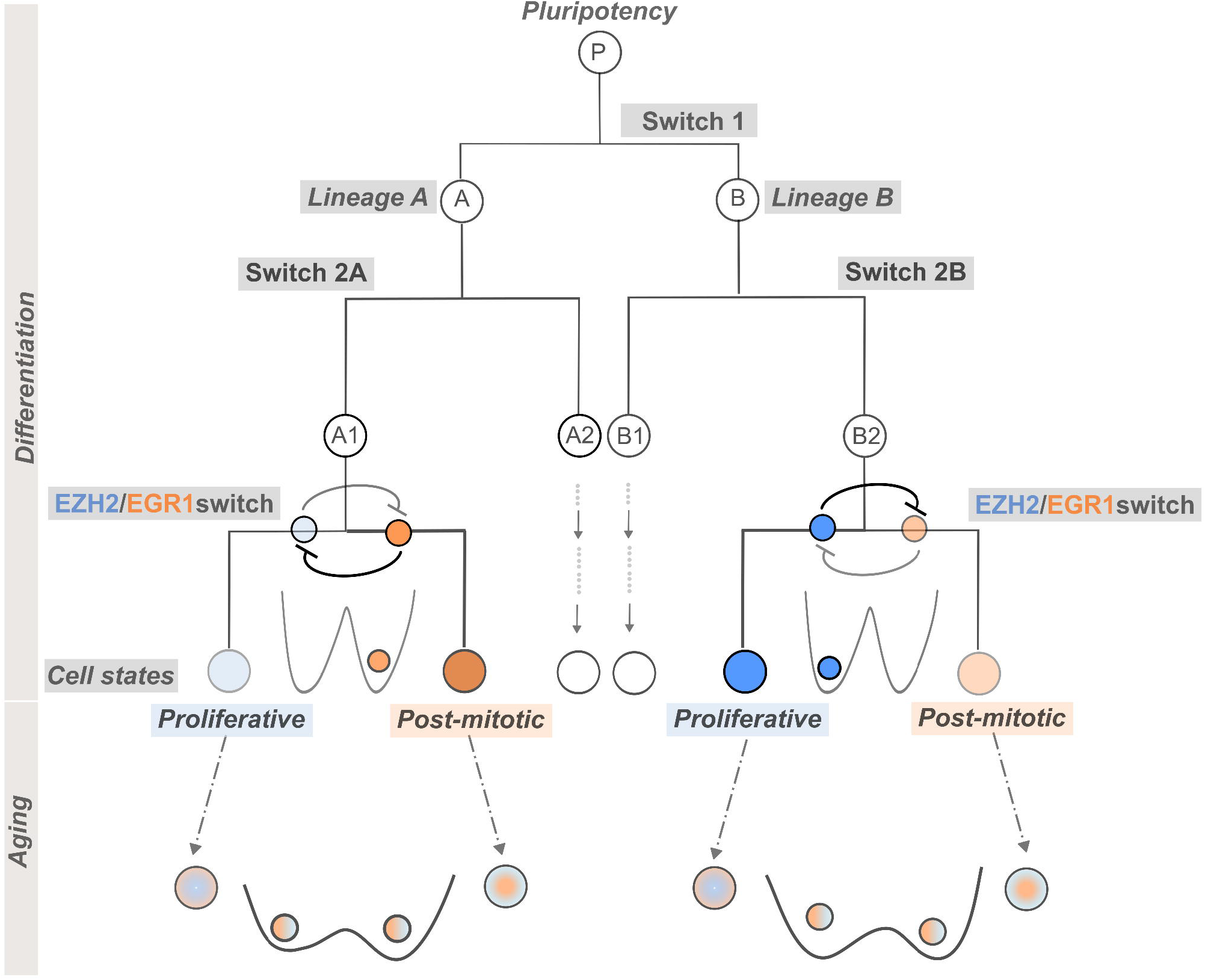
Schematic illustrating a proposed model linking development and aging through the EZH2–EGR1 genetic switch across multiple cell lineages. During development, lineage specification proceeds through a series of binary cell-fate decisions mediated by nested genetic switches. Within a given lineage, the EZH2–EGR1 switch—potentially functioning as a core module of a larger gene regulatory network—regulates the transition between proliferative progenitor states and non-proliferative, functionally specialized states. Development establishes a stable EGR1+/EZH2-transcriptional state in post-mitotic cells and an EZH2+/EGR1-state in proliferative cells. With aging, cells progressively drift away from their developmentally established state and increasingly occupy both transcriptional states, potentially reflecting a flattening of the epigenetic landscape and reduced barriers between cell states. This drift results in diminished expression of genes required for specialized cellular functions together with aberrant expression of genes that are no longer appropriate for the mature cell state, partially reversing the gene expression programs established during development. Transitions between the two states can be transiently induced in tissue specific stem cells by environmental cues, such as stress or tissue injury, to regulate the activation and proliferation.

If the EZH2-EGR1 switch is generally employed across tissues during development or tissue regeneration, we expect drift of this circuit during aging may also occurs generally, similar to what we observed in fibroblasts and in neurons. In proliferative tissues, we expect the circuit to drift from EZH2+/EGR- to EGR1+/ EZH2-states and thus may results in a similar pattern of gene expression drift across tissues. A recent study identified “mesenchymal drift” (MD) across tissues and correlated MD with tissue specific diseases of aging ^12^. MD was characterized by upregulation of mesenchymal genes and altered stromal cell composition across tissues. We found that in fibroblasts genes up-regulated in the EGR1+/EZH2-state relative to the EZH2+/EGR1-state strongly overlap with MD genes (expected overlap = 38, observed overlap=79, P-value=1.44e-11 **Figure S1C-D and Data S8**), suggesting that attenuation of the EZH2-EGR1 switch contributes significantly to MD, potentially offering a mechanistic interpretation for part of the MD signature.

Our proposed switch model of aging may have important implications for rejuvenation through transcriptional reprogramming. If attenuation of developmental genetic switches is a major driving force of aging, we should systematically search for such switches (beyond EGR1-EZH2) and target the key transcription factors involved to restore the program established by development. This model also predicts that the TFs involved in different tissues may be tissue specific, arguing that tissue specific targeting may be necessary to achieve rejuvenation and alleviate diseases.

One of the key predictions of this model is that to rejuvenate neurons, one needs to up-regulate EGR1, which is opposite to the direction of perturbation that leads to the rejuvenation of fibroblast cells (EGR1 knock down). Several lines of evidence support this prediction. Parabiosis studies in mice have shown that, in rejuvenated parabionts, EGR1 expression changes in opposite directions in proliferative cells of peripheral tissues compared with neurons in the brain. While in the proliferative cells EGR1 goes down (consistent with the direction for fibroblast rejuvenation) ^115^, in neurons EGR1 goes up ^116^, opposite to the direction of change due to aging. There is also a recent report that over-expression of EGR1 in mouse hippocampus increase dendric spine density and improve behavior in a sex dependent manner ^117^. A possible test will be to target the brain in aged mice with EGR1 over-expression to see if such intervention rejuvenates the brain and improve memory and learning.

Our model provides a framework to explain gene expression changes during aging across tissues. In a specific tissue, the switches involved should maintain expression of functional important genes and suppresses the genes not needed in that tissue. As the switches become attenuated during aging, genes not needed is de-repressed (or aberrantly expressed) and functional genes decreased. Some of the common functional genes across tissues are ribosomal genes (protein synthesis), mitochondrial genes (energy generation), and proteasome genes (protein quality control); these genes are generally down-regulated across tissues during aging. In contract, transposons and inflammation/immune response are generally not needed across tissues, and they are up-regulated during aging. The model also explains the asymmetry of down-regulated genes and up-regulated genes in term of GO enrichment. While age down-regulated genes are generally enriched for the genes functionally important for that tissue, age up-regulated genes often lack a clear functional theme ^35^ as they are related to many different alternative fates that need to be suppressed.

Our model also provides a potential mechanism for the restoration of youthful epigenetic landscape with rejuvenating TF perturbations. Sinclair and collaborators advanced the information theory of aging, which posits that repeated DNA damage and repair drives relocation of chromatin modifiers, leading to the gradual loss of youthful epigenetic information that drives aging ^18,118^. The authors hypothesized that a copy of the youthful information is stored and is read and used to re-establish the youthful epigenetic landscape during rejuvenation. With the switch model of aging, we propose that the information is encoded in the gene regulatory circuit which drifted during aging, and to rejuvenate one needs to push the circuit backwards to re-establishes the youthful transcriptional state. This is possible because the transcriptional drift is structured, e.g., in the case of passaged fibroblast is recapitulated to a large extend by a shifting between two states (**Figure 1**), even though hundreds of genes are up and down-regulated. A perturbation on a key regulator may be able to tilt the switch and shift it backwards, as long as the regulator is coupled to the larger gene regulatory network that defines the two states. Because transcription factors often function in concert with epigenetic regulators (for example, EGR1 recruits the DNA demethylase TET1 to establish locus-specific DNA methylation patterns, ^92–94^), restoring youthful transcriptional states may promote the re-establishment of DNA methylation and histone modification patterns that define a youthful epigenetic landscape ^15,119^

What could be the up-stream mechanism for the drift of the EGR1-EZH2 circuit during aging? We speculate that the loss of repression might be the upstream cause. Through the mechanism of switch, such de-repression can lead to down-regulation of functional genes. In our fibroblast perturb-seq analysis, we found that knocking down several epigenetic repressors such as the DNA methyltransferase DNMT1 accelerates aging. DNMT1 works together with PRC2 to enforce silencing/repression. Other repression mechanisms include 3D topological structure of the genome, and association with nuclear Lamins. The association with the nuclear lamina is particularly interesting, as mutations in nuclear lamins cause progeria syndromes, and lamina-associated domains (LADs) are enriched for repressive histone marks such as H3K9me3. It has been reported that during neuronal aging, there is more loss than gain of LADs ^120^. Thus, loss of LAD may be one upstream cause for the de-repression. Similarly, age-associated changes in nucleolus-associated domains (NADs) and topologically associating domains (TADs) may contribute to gene de-repression.

We note that the toggle switch circuit we constructed for fibroblasts likely only represents a small sub-circuit of a much larger gene regulatory network that defines the proliferative vs. non-proliferative state (**Figure 1**). Our previous perturb-seq screening identified dozens of single TF perturbations that either rejuvenate the aging transcriptome or accelerate aging, and most of these TF perturbations shift the relative populations of cells in the two states (**Figure 1A**, *Sengstack et. al. PNAS, 2026* ^35^). EZH2 and EGR1 are the top hits from the screening, and they function in opposite directions. There are many other TFs of which perturbation either rejuvenate or accelerate aging (such as STAT3 for immune response and ATF4 for integrated stress response), and these factors were uncovered from only a partial screen of ∼200 TFs. Thus we envision there is a much larger gene regulatory network (with which the identified TFs interact) that regulates the transition between the two states, with fine-tuned response to different cues, such as different growth factors, change of extracellular matrix stiffness and cell-cell contact, and different stresses.

Added to the complexity is that this switch must be molded by lineage specific regulators and the specific context of the cell in which it operates. For example, we showed evidence that EZH2-EGR1 circuit is involved in the transition from neural progenitor cells to matured neurons during brain development. This circuit must operate in a bigger context of gene network that regulates the transition (**Figure 5**). A well-known genetic switch downstream of Notch signaling is formed by HES1 and ASCL1 ^121^. Notch signaling induces HES1, which represses ASCL1, thereby suppressing neuronal differentiation and maintaining neural progenitor cell (NPC) proliferation. This circuit interacts with PAX6, a master regulator of neural development that establishes and patterns neural progenitors while helping orchestrate the transition from progenitor cells to differentiated neurons ^122^. Both EZH2 and HES1 are highly expressed in NPC and they cooperate to maintain NPC state ^123–125^. EZH2 also represses ASCL1 to suppress differentiation ^99,126^. Thus it is possible that a complex network involving EZH2 and EGR1 and other lineage specific factors define nested switches to orchestrate a series of developmental transitions. It will be important to reconstruct this network comprehensively to better understand the logic and design of development. Such understanding may translate into new insight into the mechanisms of aging and novel strategies for intervention.

## METHODS

### Single-cell and single nucleus RNA-seq data pre-processing

Raw gene expression count matrices underwent quality control to remove low-quality cells/nuclei, high mitochondrial-content cells, lowly expressed genes, and outliers. Donors with <200 cells/nuclei were excluded, followed by donor- or sample-wise median absolute deviation (MAD; 3 MAD) filtering. Doublets were identified with “Scrublet” (*Wolock et al., Cell Systems, 2020*) in samples containing >500 cells/nuclei and removed. Filtered count matrices were normalized to counts per million (CPM), log-transformed log 1. All analysis performed in analyzed in Scanpy. Dimensionality reduction and UMAPS: Principal component analysis (PCA) was performed using the selected highly variable genes after excluding sex-chromosome genes for samples from donors (XIST, TSIX, UTY, DDX3Y, KDM5D, EIF1AY, RPS4Y1, ZFY). The number of principal components retained was determined using elbow-plot inspection. A k-nearest neighbor (kNN) graph was constructed using the selected principal components with cosine distance metric with k optimized for each dataset, typically k ∼ 25 - 30. Uniform Manifold Approximation and Projection (UMAP) embeddings were constructed using this neighborhood graph.

### Cell-type annotation

Cell identities were assigned using canonical marker genes together with marker-gene enrichment scores computed (Scanpy’s, score_genes). Cluster-specific marker genes were identified using the Wilcoxon rank-sum test implemented in (Scanpy’s, rank_genes_groups), comparing each Leiden cluster against all remaining cells. Cell-type annotations were further validated by inspection of established marker genes.

### Differential gene expression analysis

Differential gene expression analyses were performed on log-transformed expression values using custom Python scripts implementing Welch’s t-test. Statistical significance was assessed using the Benjamini–Hochberg false discovery rate (FDR) correction for multiple tests. Genes satisfying the specified log2 fold-change (0.58, corresponding to 1.5-fold change up or down) and FDR thresholds (0.05) were considered significantly differentially expressed unless otherwise stated.

### 2D gene expression state assignment

Cells were classified into cell-states based on the detectable expression of EZH2 and EGR1 resulting in four cell states: EZH2+/EGR1+, EZH2+/EGR1−, EGR1+/EHZ2+, and EZH2−/EGR1−.

### Estimating two-dimensional density of cells in gene1-gene2 expression space

2D density of cells in gene1-gene2 (e.g., EZH2-EGR1) expression space were computed using cells expressing at least one of the two genes, excluding double-zero observations that may represent either biological absence of expression or technical dropout. Gene expression value for a single cell is given by the log transformed normalized count log 1. For density estimates, Gaussian kernels were used with Scott’s rule for bandwidth estimation (seaborn’s, kdeplot with, bw_method=”scott”), together with a empirically selected bandwidth adjustment factor (bw_adjust = 1) for visualization. Density contours were clipped to the positive expression range.

### Hierarchical clustering and heatmaps

For heatmap visualization, gene expression values were averaged within defined groups (such as donor, developmental stage, cell state) and converted to per-gene z-scores (calculated across conditions). Hierarchical clustering was performed using cosine distance with average linkage or Euclidean distance with ward linkage. Gene modules were identified by cutting the hierarchical clustering dendogram using “fcluster”, based on the visual inspection of the structure. For EZH2 and EGR1 target heatmaps, lowly expressed and low variance genes across conditions were filtered prior to clustering (quantile threshold, 0.01).

### Functional enrichment analysis

Gene ontology enrichment analyses were performed using GSEA (Python implementation, GSEAPY, with the Gene Ontology Biological Process 2025 database). Overrepresentation test and rank based gene set enrichment analysis was performed using “ClusterProfiler” in R. Selected analyses were additionally validated using PANTHER database for over-representation analysis. Gene sets with Benjamini–Hochberg adjusted P-values < 0.05 were considered significantly enriched and retained for analysis.

### Correlation analysis

All gene-pair correlations were computed using Pearson’s correlation coefficient. Unless otherwise stated, all statistical tests were two-sided, and multiple hypothesis testing was controlled using the Benjamini–Hochberg false discovery rate (FDR) procedure.

### Identification of EZH2 and EGR1 target genes

EZH2 target genes in fibroblasts were identified from previously published EZH2 ChIP-seq dataset (*Moqri, M. et al., Nature Communications, 2024*). Reproducible EZH2 binding sites were identified using the Irreproducible Discovery Rate (IDR) framework (IDR < 0.05), and consensus peaks were assigned to genes based on overlap with promoter regions (2.5 kb) from transcription start sites) using GENCODE annotations. EZH2 occupancy was quantified from the corresponding merged fold-enrichment bigWig files. The union of neonatal and aged promoter-associated genes was used as the final EZH2 target gene set (∼2,280 genes). Neuronal EZH2 target genes were defined as the intersection of fibroblast EZH2 target genes and genes marked by H3K27me3 in human neural progenitor cells and neurons during gestation, as identified by CUT&Tag profiling (*Ditzer et. al, Neuron 2025)*.

EGR1 target genes. Predicted EGR1 target genes were obtained from the GSEA transcription factor target gene set (*Xie et al., 2005; Subramanian et al., 2005; Liberzon et al., 2011),* based on the occurrence of the EGR1-binding motifs within 2 kb of the transcription start site (TSS). Additionally, an independent motif-based prediction was performed by scanning (-5kb of TSS) for EGR1 binding motifs. The predicted genes were intersected with genes that satisfy the Gene Ontology (GO) annotations corresponding to neuronal processes. The intersection of these GO-derived gene sets with the predicted EGR1 target genes were then defined as neuron specific-EGR1 targets.

### Fibroblast CRISPR perturbation analyses

Previously published CRISPR activation/interference (CRISPRa/i) perturbation data from human fibroblasts were obtained from (*Sengstack, et. al. PNAS 2026)*. Raw count matrices were processed, normalized and log-transformed prior to downstream analyses, and differential gene expression was computed for each perturbation relative to non-targeting control cells as described. Differential expression vectors for each perturbation were compared with differential expression observed during replicative aging (PD32 versus PD14 fibroblasts) using Pearson correlation as described in Sengstack et al. PNAS 2026. Perturbations exhibiting significant positive correlations (r 0.35) were classified as aging-accelerating perturbations, whereas perturbations exhibiting significant negative correlations (r -0.35) were classified as rejuvenating perturbations.

Marker genes were selected from genes differentially expressed during replicative aging together with genes with established functional roles in cell proliferation and extracellular matrix maintenance.

### RNA velocity analyses

Spliced and unspliced RNA counts were obtained from raw counts using velocyto *(La Manno et. al., Nature 2018).* Data filtering, processing, HVG selection (∼3000 genes) was performed as described in the earlier sections of Methods. UMAP projection was done with PCs 30 and kNN=30 and HVG= 3000. RNA velocity vectors were inferred from fits to the full “dynamical model” in scVelo (*Bergen et. al. 2020*, Nature Biotechnology) and projected onto the UMAP embedding.

### Human Neuronal development single-nucleus RNA

Single-nucleus RNA sequencing data during development was obtained from the raw files (*Velmeshev et. al. 2023*). Analyses were restricted to neocortical regions annotated as PFC, BA9/46, BA10, BA8, STG, BA22, temporal cortex, primary motor cortex, frontoparietal cortex, frontal cortex and cortex. Cell-type identities as annotated in the original dataset were retained. UMAPs were constructed using the top 4,000 HVGs across developmental stages, with Jaccard similarity as the distance metric, and parameters PCs =15 and kNN =25.

### Pharmacological inhibition of EZH2 during cortical neuronal maturation from iPSCs

All the bulk RNA-sequencing data were processed using PyDESEQ2 framework *(Muzellec et. al. Bioinformatics 2023)*. Published bulk RNA-sequencing data (GEO: GEO: GSE1226223) were downloaded from and analyzed using the PyDESeq2 package. In the original experiments, neural stem cells were treated with GSK343, a selective pharmacological inhibitor of EZH2, from day 12 to day 20 of differentiation. Differential gene expression analysis was performed on day 35, when the iPSC-derived cells had matured into cortical neurons, to assess the transcriptional consequences of transient EZH2 inhibition during neuronal differentiation *(Ciceri, G., et. al. Nature 2024)*.

### Neuronal aging: single-nucleus RNA-seq data of dorsolateral prefrontal cortex

Single-nucleus RNA-sequencing (snRNA-seq) raw count data spanning the human lifespan were obtained from the published dataset by *Catching et al., Cell Rep. 2026*. To minimize inter-donor variability, analyses were primarily restricted to the NABEC cohort (individuals of European ancestry) and female samples. Excitatory neurons were selected as per the annotation in the dataset. Cell-annotation and labeling was performed to identify the excitatory neuron types using the gene markers from the Velmeshev’s dataset as described in the section “cell-type annotation”. Subsequent analyses focused on female donors and upper-layer (L2–L3) excitatory neurons. Donors aged 10–30 years were classified as the young group, whereas donors older than 80 years were classified as the aged group.

Pairwise gene-gene correlation matrix for top 30 genes positively correlated with EGR1 and top 30 genes positively correlated with EZH2 were computed from donor-level pseudobulk expression profiles across all female donors (n=58) spanning 10–100 years of age.

Aging-associated target gene heatmaps: heatmaps were generated using the 65 EGR1 target genes identified from the developmental module (Cluster 1) exhibiting increasing expression during maturation (Figure 2D).

### Human single-cell hematopoietic and muscle stem aging dataset

The published dataset for human hematopoietic stem cell single-cell expression was re-analyzed (*Li et al. Nature Methods 2025*). The study quantified the gene expression from prenatal liver samples, up to donors aged > 70 years. In this study, only the cells from hematopoietic stem cell compartment were considered. The developmental stage and cell-type annotations provided in the original studies were retained for all analyses.

The published processed human skeletal muscle single-cell transcriptome atlas (*Lai et al., Nature 2024*) was re-analyzed. The study profiled hindlimb skeletal muscle biopsies (semitendinosus, vastus lateralis, and gluteus medius) obtained from healthy adult (15–46 years, n = 12) and older (74–99 years, n = 19) donors using single-cell and single-nucleus RNA sequencing. We analyzed only the muscle stem cell populations from the annotated data, retaining the classification for different cell states i.e. (quiescent, early primed, late primed and differentiated myogenic cells). 2D density plots were constructed as described.

**Table 1:**
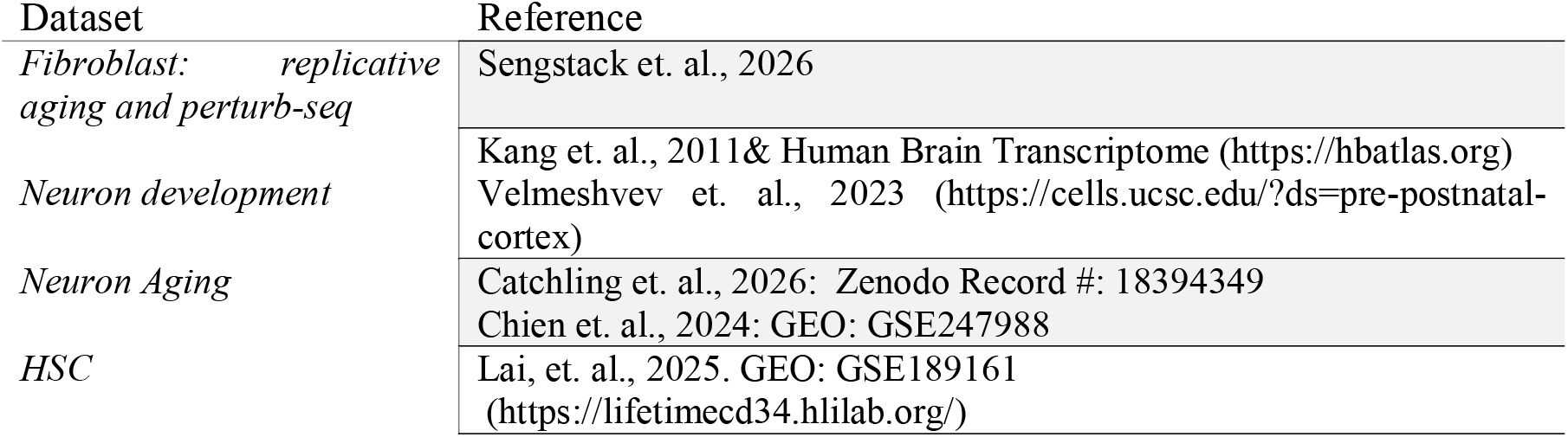
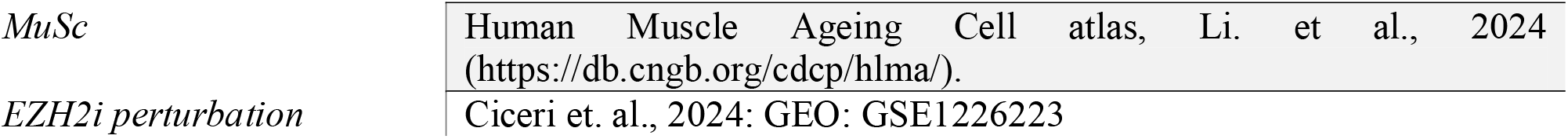
Summary of datasets and download links in this study.

## Data and Code availability

will be made available at the time of publication

## Acknowledgements

We thank Drs. Elizabeth Blackburn and Peter Walter for helpful discussions.

## Author Contributions

NK and HL conceived and designed the study. NK performed computational/bioinformatic analyses. JZ and EW assisted in bioinformatic analyses. CD contributed to fibroblast experiments. NK and HL analyzed and interpreted data. NK and HL wrote the manuscript.

## References

1 Kenyon, C., Chang, J., Gensch, E., Rudner, A. & Tabtiang, R. A C. elegans mutant that lives twice as long as wild type. Nature 366, 461–464 (1993). 10.1038/366461a0

2 Lin, K., Dorman, J. B., Rodan, A. & Kenyon, C. daf-16: An HNF-3/forkhead family member that can function to double the life-span of Caenorhabditis elegans. Science 278, 1319–1322 (1997). 10.1126/science.278.5341.1319

3 Kimura, K. D., Tissenbaum, H. A., Liu, Y. & Ruvkun, G. daf-2, an insulin receptor-like gene that regulates longevity and diapause in Caenorhabditis elegans. Science 277, 942–946 (1997). 10.1126/science.277.5328.942

4 Ogg, S. et al. The Fork head transcription factor DAF-16 transduces insulin-like metabolic and longevity signals in C. elegans. Nature 389, 994–999 (1997). 10.1038/40194

5 Walker, R. F. A Mechanistic Theory of Development-Aging Continuity in Humans and Other Mammals. Cells 11 (2022). 10.3390/cells11050917

6 Walker, R. F. Developmental theory of aging revisited: focus on causal and mechanistic links between development and senescence. Rejuvenation Res 14, 429– 436 (2011). 10.1089/rej.2011.1162

7 de Magalhaes, J. P. Programmatic features of aging originating in development: aging mechanisms beyond molecular damage? FASEB J 26, 4821–4826 (2012). 10.1096/fj.12-210872

8 de Magalhaes, J. P. An overview of contemporary theories of ageing. Nat Cell Biol 27, 1074–1082 (2025). 10.1038/s41556-025-01698-7

9 Blagosklonny, M. V. Aging and immortality: quasi-programmed senescence and its pharmacologic inhibition. Cell Cycle 5, 2087–2102 (2006). 10.4161/cc.5.18.3288

10 Blagosklonny, M. V. MTOR-driven quasi-programmed aging as a disposable soma theory: blind watchmaker vs. intelligent designer. Cell Cycle 12, 1842–1847 (2013). 10.4161/cc.25062

11 Gems, D. The hyperfunction theory: An emerging paradigm for the biology of aging. Ageing Res Rev 74, 101557 (2022). 10.1016/j.arr.2021.101557

12 Lu, J. Y. et al. Prevalent mesenchymal drift in aging and disease is reversed by partial reprogramming. Cell 188, 5895–5911 e5817 (2025). 10.1016/j.cell.2025.07.031

13 Connolly, E. et al. Loss of immune cell identity with age inferred from large atlases of single cell transcriptomes. Aging Cell 23, e14306 (2024). 10.1111/acel.14306

14 Salzer, M. C. et al. Identity Noise and Adipogenic Traits Characterize Dermal Fibroblast Aging. Cell 175, 1575–1590 e1522 (2018). 10.1016/j.cell.2018.10.012

15 Yucel, A. D. & Gladyshev, V. N. Systemic epigenetic dysregulation as a driver of ageing and a therapeutic target. Nat Rev Mol Cell Biol 27, 528–542 (2026). 10.1038/s41580-026-00958-0

16 Gorelov, R. & Hochedlinger, K. A cellular identity crisis? Plasticity changes during aging and rejuvenation. Genes Dev 38, 823–842 (2024). 10.1101/gad.351728.124

17 Gruber, J., Yee, Z. & Tolwinski, N. S. Developmental Drift and the Role of Wnt Signaling in Aging. Cancers (Basel) 8 (2016). 10.3390/cancers8080073

18 Lu, Y. R., Tian, X. & Sinclair, D. A. The Information Theory of Aging. Nat Aging 3, 1486–1499 (2023). 10.1038/s43587-023-00527-6

19. Horvath, S. DNA methylation age of human tissues and cell types. Genome Biol 14, R115 (2013). 10.1186/gb-2013-14-10-r115

20 Lu, A. T. et al. Universal DNA methylation age across mammalian tissues. Nat Aging 3, 1144–1166 (2023). 10.1038/s43587-023-00462-6

21 Moqri, M. et al. PRC2-AgeIndex as a universal biomarker of aging and rejuvenation. Nat Commun 15, 5956 (2024). 10.1038/s41467-024-50098-2

22 Tyshkovskiy, A. et al. Universal transcriptomic hallmarks of mammalian ageing and mortality. Nature 654, 173–188 (2026). 10.1038/s41586-026-10542-3

23 de Magalhaes, J. P., Curado, J. & Church, G. M. Meta-analysis of age-related gene expression profiles identifies common signatures of aging. Bioinformatics 25, 875– 881 (2009). 10.1093/bioinformatics/btp073

24 Palmer, D., Fabris, F., Doherty, A., Freitas, A. A. & de Magalhaes, J. P. Ageing transcriptome meta-analysis reveals similarities and differences between key mammalian tissues. Aging (Albany NY) 13, 3313–3341 (2021). 10.18632/aging.202648

25 Lu, Z. et al. Organism-wide cellular dynamics and epigenomic remodeling in mammalian aging. Science 391, eadw6273 (2026). 10.1126/science.adw6273

26 Matsuzaki, T., Weistuch, C., de Graff, A., Dill, K. A. & Balazsi, G. Transcriptional drift in aging cells: A global decontroller. Proc Natl Acad Sci U S A 121, e2401830121 (2024). 10.1073/pnas.2401830121

27 Colantuoni, C. et al. Temporal dynamics and genetic control of transcription in the human prefrontal cortex. Nature 478, 519–523 (2011). 10.1038/nature10524

28 Donertas, H. M. et al. Gene expression reversal toward pre-adult levels in the aging human brain and age-related loss of cellular identity. Sci Rep 7, 5894 (2017). 10.1038/s41598-017-05927-4

29 Somel, M., et al. MicroRNA, mRNA, and protein expression link development and aging in human and macaque brain. Genome Res 20, 1207–1218 (2010). 10.1101/gr.106849.110

30 Izgi, H. et al. Inter-tissue convergence of gene expression during ageing suggests age-related loss of tissue and cellular identity. Elife 11 (2022). 10.7554/eLife.68048

31 Lu, Y. et al. Reprogramming to recover youthful epigenetic information and restore vision. Nature 588, 124–129 (2020). 10.1038/s41586-020-2975-4

32 Ocampo, A. et al. In Vivo Amelioration of Age-Associated Hallmarks by Partial Reprogramming. Cell 167, 1719–1733 e1712 (2016). 10.1016/j.cell.2016.11.052

33 Browder, K. C. et al. In vivo partial reprogramming alters age-associated molecular changes during physiological aging in mice. Nat Aging 2, 243–253 (2022). 10.1038/s43587-022-00183-2

34 Sarkar, T. J. et al. Transient non-integrative expression of nuclear reprogramming factors promotes multifaceted amelioration of aging in human cells. Nat Commun 11, 1545 (2020). 10.1038/s41467-020-15174-3

35 Sengstack, J. et al. Systematic identification of single transcription factor perturbations that drive cellular and tissue rejuvenation. Proc Natl Acad Sci U S A 123, e2515183123 (2026). 10.1073/pnas.2515183123

36 Ribeiro, R. et al. In vivo cyclic induction of the FOXM1 transcription factor delays natural and progeroid aging phenotypes and extends healthspan. Nat Aging 2, 397– 411 (2022). 10.1038/s43587-022-00209-9

37 Ferreira, F. J. et al. FOXM1 expression reverts aging chromatin profiles through repression of the senescence-associated pioneer factor AP-1. Nat Commun 16, 2931 (2025). 10.1038/s41467-025-57503-4

38 Ping, J. et al. Human immune aging clock identifies RUNX1 as a decelerator of T cell senescence. Immunity 59, 1039–1057 e1011 (2026). 10.1016/j.immuni.2026.02.007

39 Zhou, C. et al. Runx1 protects against the pathological progression of osteoarthritis. Bone Res 9, 50 (2021). 10.1038/s41413-021-00173-x

40 Jing, Y. et al. Genome-wide CRISPR activation screening in senescent cells reveals SOX5 as a driver and therapeutic target of rejuvenation. Cell Stem Cell 30, 1452– 1471 e1410 (2023). 10.1016/j.stem.2023.09.007

41 Wang, W. et al. A genome-wide CRISPR-based screen identifies KAT7 as a driver of cellular senescence. Sci Transl Med 13 (2021). 10.1126/scitranslmed.abd2655

42 Korotkov, A., Seluanov, A. & Gorbunova, V. Sirtuin 6: linking longevity with genome and epigenome stability. Trends Cell Biol 31, 994–1006 (2021). 10.1016/j.tcb.2021.06.009

43 Nagar, R. et al. SIRT6 overexpression counteracts chromatin aging in the male murine liver. Nat Commun 17 (2026). 10.1038/s41467-026-73115-y

44 Hayflick, L. & Moorhead, P. S. The serial cultivation of human diploid cell strains. Exp Cell Res 25, 585–621 (1961). 10.1016/0014-4827(61)90192-6

45 Ito, T., Teo, Y. V., Evans, S. A., Neretti, N. & Sedivy, J. M. Regulation of Cellular Senescence by Polycomb Chromatin Modifiers through Distinct DNA Damage- and Histone Methylation-Dependent Pathways. Cell Rep 22, 3480–3492 (2018). 10.1016/j.celrep.2018.03.002

46 Yang, N. et al. A hyper-quiescent chromatin state formed during aging is reversed by regeneration. Mol Cell 83, 1659–1676 e1611 (2023). 10.1016/j.molcel.2023.04.005

47 Chen, H. et al. Polycomb protein Ezh2 regulates pancreatic beta-cell Ink4a/Arf expression and regeneration in diabetes mellitus. Genes Dev 23, 975–985 (2009). 10.1101/gad.1742509

48 Zhang, Y., Yu, C., Agborbesong, E. & Li, X. Downregulation of EZH2 Promotes Renal Epithelial Cellular Senescence and Kidney Aging. FASEB J 39, e70605 (2025). 10.1096/fj.202500128R

49 Ezhkova, E. et al. Ezh2 orchestrates gene expression for the stepwise differentiation of tissue-specific stem cells. Cell 136, 1122–1135 (2009). 10.1016/j.cell.2008.12.043

50 Li, Y. et al. Ezh2 Inhibits Replicative Senescence of Atrial Fibroblasts Through Promotion of H3K27me3 in the Promoter Regions of CDKN2a and Timp4 Genes. J Inflamm Res 15, 4693–4708 (2022). 10.2147/JIR.S374951

51 Levy, D. E. & Darnell, J. E., Jr. Stats: transcriptional control and biological impact. Nat Rev Mol Cell Biol 3, 651–662 (2002). 10.1038/nrm909

52 Harding, H. P. et al. An integrated stress response regulates amino acid metabolism and resistance to oxidative stress. Mol Cell 11, 619–633 (2003). 10.1016/s1097-2765(03)00105-9

53 Costa-Mattioli, M. & Walter, P. The integrated stress response: From mechanism to disease. Science 368 (2020). 10.1126/science.aat5314

54 Baron, V., Adamson, E. D., Calogero, A., Ragona, G. & Mercola, D. The transcription factor Egr1 is a direct regulator of multiple tumor suppressors including TGFbeta1, PTEN, p53, and fibronectin. Cancer Gene Ther 13, 115–124 (2006). 10.1038/sj.cgt.7700896

55 Bhattacharyya, S., Fang, F., Tourtellotte, W. & Varga, J. Egr-1: new conductor for the tissue repair orchestra directs harmony (regeneration) or cacophony (fibrosis). J Pathol 229, 286–297 (2013). 10.1002/path.4131

56 Gashler, A. & Sukhatme, V. P. Early growth response protein 1 (Egr-1): prototype of a zinc-finger family of transcription factors. Prog Nucleic Acid Res Mol Biol 50, 191–224 (1995). 10.1016/s0079-6603(08)60815-6

57 Bhattacharyya, S. et al. Early growth response transcription factors: key mediators of fibrosis and novel targets for anti-fibrotic therapy. Matrix Biol 30, 235–242 (2011). 10.1016/j.matbio.2011.03.005

58 Bhattacharyya, S. et al. Egr-1 induces a profibrotic injury/repair gene program associated with systemic sclerosis. PLoS One 6, e23082 (2011). 10.1371/journal.pone.0023082

59 Krones-Herzig, A., Adamson, E. & Mercola, D. Early growth response 1 protein, an upstream gatekeeper of the p53 tumor suppressor, controls replicative senescence. Proc Natl Acad Sci U S A 100, 3233–3238 (2003). 10.1073/pnas.2628034100

60 Rognoni, E. et al. Fibroblast state switching orchestrates dermal maturation and wound healing. Mol Syst Biol 14, e8174 (2018). 10.15252/msb.20178174

61 Bracken, A. P. et al. The Polycomb group proteins bind throughout the INK4A-ARF locus and are disassociated in senescent cells. Genes Dev 21, 525–530 (2007). 10.1101/gad.415507

62 Pilling, L. C. et al. Human longevity: 25 genetic loci associated in 389,166 UK biobank participants. Aging (Albany NY) 9, 2504–2520 (2017). 10.18632/aging.101334

63 Bracken, A. P. et al. EZH2 is downstream of the pRB-E2F pathway, essential for proliferation and amplified in cancer. EMBO J 22, 5323–5335 (2003). 10.1093/emboj/cdg542

64 Chen, S. J. et al. The early-immediate gene EGR-1 is induced by transforming growth factor-beta and mediates stimulation of collagen gene expression. J Biol Chem 281, 21183–21197 (2006). 10.1074/jbc.M603270200

65 Liberzon, A. et al. Molecular signatures database (MSigDB) 3.0. Bioinformatics 27, 1739–1740 (2011). 10.1093/bioinformatics/btr260

66 Guan, X. et al. EZH2 overexpression dampens tumor-suppressive signals via an EGR1 silencer to drive breast tumorigenesis. Oncogene 39, 7127–7141 (2020). 10.1038/s41388-020-01484-9

67 Tanaka, S. et al. Ezh2 augments leukemogenicity by reinforcing differentiation blockage in acute myeloid leukemia. Blood 120, 1107–1117 (2012). 10.1182/blood-2011-11-394932

68 Ditzer, N. et al. Epigenome profiling identifies H3K27me3 regulation of extracellular matrix composition in human corticogenesis. Neuron 113, 2927–2944. e2910 (2025).

69 Neo, W. H. et al. MicroRNA miR-124 controls the choice between neuronal and astrocyte differentiation by fine-tuning Ezh2 expression. J Biol Chem 289, 20788– 20801 (2014). 10.1074/jbc.M113.525493

70 Lee, S. W., Oh, Y. M., Lu, Y. L., Kim, W. K. & Yoo, A. S. MicroRNAs Overcome Cell Fate Barrier by Reducing EZH2-Controlled REST Stability during Neuronal Conversion of Human Adult Fibroblasts. Dev Cell 46, 73–84 e77 (2018). 10.1016/j.devcel.2018.06.007

71 Guajardo, L. et al. Downregulation of the Polycomb-Associated Methyltransferase Ezh2 during Maturation of Hippocampal Neurons Is Mediated by MicroRNAs Let-7 and miR-124. Int J Mol Sci 21 (2020). 10.3390/ijms21228472

72 Ptashne, M. Lambda’s switch: lessons from a module swap. Curr Biol 16, R459–462 (2006). 10.1016/j.cub.2006.05.037

73 Acar, M., Becskei, A. & van Oudenaarden, A. Enhancement of cellular memory by reducing stochastic transitions. Nature 435, 228–232 (2005). 10.1038/nature03524

74 Gardner, T. S., Cantor, C. R. & Collins, J. J. Construction of a genetic toggle switch in Escherichia coli. Nature 403, 339–342 (2000). 10.1038/35002131

75 Huang, S., Guo, Y. P., May, G. & Enver, T. Bifurcation dynamics in lineage-commitment in bipotent progenitor cells. Dev Biol 305, 695–713 (2007). 10.1016/j.ydbio.2007.02.036

76 Imayoshi, I. et al. Oscillatory control of factors determining multipotency and fate in mouse neural progenitors. Science 342, 1203–1208 (2013). 10.1126/science.1242366

77 Elowitz, M. B. & Leibler, S. A synthetic oscillatory network of transcriptional regulators. Nature 403, 335–338 (2000). 10.1038/35002125

78 Bergen, V., Lange, M., Peidli, S., Wolf, F. A. & Theis, F. J. Generalizing RNA velocity to transient cell states through dynamical modeling. Nat Biotechnol 38, 1408–1414 (2020). 10.1038/s41587-020-0591-3

79 Herring, C. A. et al. Human prefrontal cortex gene regulatory dynamics from gestation to adulthood at single-cell resolution. Cell 185, 4428–4447 e4428 (2022). 10.1016/j.cell.2022.09.039

80 Ramos, S. I. et al. An atlas of late prenatal human neurodevelopment resolved by single-nucleus transcriptomics. Nat Commun 13, 7671 (2022). 10.1038/s41467-022-34975-2

81 Trevino, A. E. et al. Chromatin and gene-regulatory dynamics of the developing human cerebral cortex at single-cell resolution. Cell 184, 5053–5069 e5023 (2021). 10.1016/j.cell.2021.07.039

82 Velmeshev, D. et al. Single-cell analysis of prenatal and postnatal human cortical development. Science 382, eadf0834 (2023). 10.1126/science.adf0834

83 Kang, H. J. et al. Spatio-temporal transcriptome of the human brain. Nature 478, 483–489 (2011). 10.1038/nature10523

84 Nowakowski, T. J. et al. Spatiotemporal gene expression trajectories reveal developmental hierarchies of the human cortex. Science 358, 1318–1323 (2017). 10.1126/science.aap8809

85 Bystron, I., Blakemore, C. & Rakic, P. Development of the human cerebral cortex: Boulder Committee revisited. Nat Rev Neurosci 9, 110–122 (2008). 10.1038/nrn2252

86 Bayatti, N. et al. A molecular neuroanatomical study of the developing human neocortex from 8 to 17 postconceptional weeks revealing the early differentiation of the subplate and subventricular zone. Cereb Cortex 18, 1536–1548 (2008). 10.1093/cercor/bhm184

87 Silbereis, J. C., Pochareddy, S., Zhu, Y., Li, M. & Sestan, N. The Cellular and Molecular Landscapes of the Developing Human Central Nervous System. Neuron 89, 248–268 (2016). 10.1016/j.neuron.2015.12.008

88 Moqri, M. et al. PRC2-AgeIndex as a universal biomarker of aging and rejuvenation. Nature Communications 15, 5956 (2024).

89 Ditzer, N. et al. Epigenome profiling identifies H3K27me3 regulation of extracellular matrix composition in human corticogenesis. Neuron 113, 2927–2944 e2910 (2025). 10.1016/j.neuron.2025.06.016

90 Ciceri, G. et al. An epigenetic barrier sets the timing of human neuronal maturation. Nature 626, 881–890 (2024). 10.1038/s41586-023-06984-8

91 Purzner, J. et al. Ezh2 Delays Activation of Differentiation Genes During Normal Cerebellar Granule Neuron Development and in Medulloblastoma. bioRxiv (2024). 10.1101/2024.11.21.624171

92 Chien, J. F. et al. Cell-type-specific effects of age and sex on human cortical neurons. Neuron 112, 2524–2539 e2525 (2024). 10.1016/j.neuron.2024.05.013

93 Sun, Z. et al. EGR1 recruits TET1 to shape the brain methylome during development and upon neuronal activity. Nat Commun 10, 3892 (2019). 10.1038/s41467-019-11905-3

94 Rudenko, A. et al. Tet1 is critical for neuronal activity-regulated gene expression and memory extinction. Neuron 79, 1109–1122 (2013). 10.1016/j.neuron.2013.08.003

95 Zhang, R. R. et al. Tet1 regulates adult hippocampal neurogenesis and cognition. Cell Stem Cell 13, 237–245 (2013). 10.1016/j.stem.2013.05.006

96 Penner, M. R. et al. Age-related changes in Egr1 transcription and DNA methylation within the hippocampus. Hippocampus 26, 1008–1020 (2016). 10.1002/hipo.22583

97 Jeffries, A. M. et al. Single-cell transcriptomic and genomic changes in the ageing human brain. Nature 646, 657–666 (2025). 10.1038/s41586-025-09435-8

98 Catching, A. et al. Single-nucleus multiome analysis in the human prefrontal cortex identifies gene expression and cis-regulatory elements associated with aging. Cell Rep 45, 117110 (2026). 10.1016/j.celrep.2026.117110

99 Pereira, J. D. et al. Ezh2, the histone methyltransferase of PRC2, regulates the balance between self-renewal and differentiation in the cerebral cortex. Proc Natl Acad Sci U S A 107, 15957–15962 (2010). 10.1073/pnas.1002530107

100 Aoki, R. et al. The polycomb group gene product Ezh2 regulates proliferation and differentiation of murine hepatic stem/progenitor cells. J Hepatol 52, 854–863 (2010). 10.1016/j.jhep.2010.01.027

101 Xie, H. et al. Polycomb repressive complex 2 regulates normal hematopoietic stem cell function in a developmental-stage-specific manner. Cell Stem Cell 14, 68–80 (2014). 10.1016/j.stem.2013.10.001

102 Kulkarni, R. Early Growth Response Factor 1 in Aging Hematopoietic Stem Cells and Leukemia. Front Cell Dev Biol 10, 925761 (2022). 10.3389/fcell.2022.925761

103 Min, I. M. et al. The transcription factor EGR1 controls both the proliferation and localization of hematopoietic stem cells. Cell Stem Cell 2, 380–391 (2008). 10.1016/j.stem.2008.01.015

104 Desterke, C., Bennaceur-Griscelli, A. & Turhan, A. G. EGR1 dysregulation defines an inflammatory and leukemic program in cell trajectory of human-aged hematopoietic stem cells (HSC). Stem Cell Res Ther 12, 419 (2021). 10.1186/s13287-021-02498-0

105 Li, H. et al. The dynamics of hematopoiesis over the human lifespan. Nat Methods 22, 422–434 (2025). 10.1038/s41592-024-02495-0

106 Kowalczyk, M. S. et al. Single-cell RNA-seq reveals changes in cell cycle and differentiation programs upon aging of hematopoietic stem cells. Genome Res 25, 1860–1872 (2015). 10.1101/gr.192237.115

107 Muto, T. et al. Concurrent loss of Ezh2 and Tet2 cooperates in the pathogenesis of myelodysplastic disorders. J Exp Med 210, 2627–2639 (2013). 10.1084/jem.20131144

108 Pan, S. et al. PURE-seq identifies Egr1 as a Potential Master Regulator in Murine Aging by Sequencing Long-Term Hematopoietic Stem Cells. bioRxiv (2024). 10.1101/2024.08.12.607664

109 Singh, S., Jakubison, B. & Keller, J. R. Protection of hematopoietic stem cells from stress-induced exhaustion and aging. Curr Opin Hematol 27, 225–231 (2020). 10.1097/MOH.0000000000000586

110 Janzen, V. et al. Stem-cell ageing modified by the cyclin-dependent kinase inhibitor p16INK4a. Nature 443, 421–426 (2006). 10.1038/nature05159

111 Flach, J. et al. Replication stress is a potent driver of functional decline in ageing haematopoietic stem cells. Nature 512, 198–202 (2014). 10.1038/nature13619

112 Pang, W. W. et al. Human bone marrow hematopoietic stem cells are increased in frequency and myeloid-biased with age. Proc Natl Acad Sci U S A 108, 20012–20017 (2011). 10.1073/pnas.1116110108

113 Lai, Y. et al. Multimodal cell atlas of the ageing human skeletal muscle. Nature 629, 154–164 (2024). 10.1038/s41586-024-07348-6

114 Zhang, W. et al. Transcription factor EGR1 promotes differentiation of bovine skeletal muscle satellite cells by regulating MyoG gene expression. J Cell Physiol 233, 350–362 (2018). 10.1002/jcp.25883

115 Palovics, R. et al. Molecular hallmarks of heterochronic parabiosis at single-cell resolution. Nature 603, 309–314 (2022). 10.1038/s41586-022-04461-2

116 Villeda, S. A. et al. Young blood reverses age-related impairments in cognitive function and synaptic plasticity in mice. Nat Med 20, 659–663 (2014). 10.1038/nm.3569

117 Rocks, D. et al. Egr1 is a sex-specific regulator of neuronal chromatin, synaptic plasticity, and behaviour. bioRxiv (2023). 10.1101/2023.12.20.572697

118 Yang, J. H. et al. Loss of epigenetic information as a cause of mammalian aging. Cell 186, 305–326 e327 (2023). 10.1016/j.cell.2022.12.027

119 Zhu, M. et al. Reduction of DNA Topoisomerase Top2 Reprograms the Epigenetic Landscape and Extends Health and Life Span Across Species. Aging Cell 24, e70010 (2025). 10.1111/acel.70010

120 Zhang, C. et al. Substructure and maturation of lamina-associated domains in neurons of the developing and adult human brain. bioRxiv (2025). 10.1101/2024.11.27.625786

121 Sueda, R., Imayoshi, I., Harima, Y. & Kageyama, R. High Hes1 expression and resultant Ascl1 suppression regulate quiescent vs. active neural stem cells in the adult mouse brain. Genes Dev 33, 511–523 (2019). 10.1101/gad.323196.118

122 Sansom, S. N. et al. The level of the transcription factor Pax6 is essential for controlling the balance between neural stem cell self-renewal and neurogenesis. PLoS Genet 5, e1000511 (2009). 10.1371/journal.pgen.1000511

123 Hirabayashi, Y. et al. Polycomb limits the neurogenic competence of neural precursor cells to promote astrogenic fate transition. Neuron 63, 600–613 (2009). 10.1016/j.neuron.2009.08.021

124 Kageyama, R., Ohtsuka, T. & Kobayashi, T. The Hes gene family: repressors and oscillators that orchestrate embryogenesis. Development 134, 1243–1251 (2007). 10.1242/dev.000786

125 Kunoh, S., Nakashima, H. & Nakashima, K. Epigenetic Regulation of Neural Stem Cells in Developmental and Adult Stages. Epigenomes 8 (2024). 10.3390/epigenomes8020022

126 Sher, F., Boddeke, E., Olah, M. & Copray, S. Dynamic changes in Ezh2 gene occupancy underlie its involvement in neural stem cell self-renewal and differentiation towards oligodendrocytes. PLoS One 7, e40399 (2012). 10.1371/journal.pone.0040399

