## Supplementary Information for "A genetic switch involving EGR1 and EZH2 links aging and development across human tissues"

Supplementary material include:

1. Supplementary Data Tables (S1 – S14)

2. Supplementary Figures (Figure S1 - Figure S3)

**1. Supplementary Data Tables:**

1. Data S1: Log2fc expression values of marker genes in CRA or CRI perturbations relatives to non-targeting control for rejuvenating and age-accelerating perturbations as in Fig1A.
2. Data S2: Predicted EGR1 target genes.
3. Data S3: Overrepresentation analysis (ORA) of predicted EGR1 targets (PANTHER).
4. Data S4: EZH2 target genes in fibroblasts and neurons.
5. Data S5: Overrepresentation analysis (ORA) of EZH2 target genes (PANTHER)
6. Data S6: DEG between two states i.e. EGR1+/EZH2- vs. EZH2+/EGR1- in PD32 and PD14.
7. Data S7: Gene Set Enrichment Analysis (GSEA) of genes in EGR1+/EZH2- vs. EZH2+/EGR1- in PD32.
8. Data S8: Mesenchymal Drift (“MD”) genes overlap with genes up-regulated in the EGR1+/EZH2- state relative to the EZH2+/EGR1- state in PD 32 fibroblast cells.
9. Data S9: Overrepresentation analysis (ORA) of EGR1 targets during neuronal development
10. Data S10: Overrepresentation analysis (ORA) of EZH2 targets during neuronal development.
11. Data S11: Overrepresentation analysis (ORA) of genes overlapping between those induced by EZH2 inhibition during in vitro maturation of neurons and EGR1 neuronal targets
12. Data S12: Overrepresentation analysis (ORA) of genes overlapping between those induced by EZH2 inhibition during in vitro maturation of neurons and those induced by EGR1 over-expression in fibroblasts
13. Data S13: DEG between two states i.e. EZH2+/ EGR1- vs. EGR1+/EZH2- in L2/L3 neurons of female young and aged donors.
14. Data S14: Gene Set Enrichment Analysis (GSEA) of DEGs between EZH2+/ EGR1- and EGR1+/EZH2- states in L2/L3 neurons of female young and aged donors

#### 2. Supplementary Figures

**Figure S1: EZH2-EGR1 regulatory network in fibroblasts**. **A)** Correlation between EZH2 expression and marker genes across perturbations (as in Figure 1A) in PD32 fibroblasts. Green: rejuvenating perturbations, black: aging-accelerating perturbations. **B)** Overlap between EGR1-responsive genes (Bhattacharrya et. al. 2011, PLoS One) and genes up-regulated in the EGR1+/EZH2- state relative to the EZH2+/EGR1- state in PD 32 fibroblast cells. **C)** Overlap between mesenchymal drift genes (MD genes) from ( Lu. et. al., Cell, 2025 ) and genes up-regulated in the EGR1+/EZH2- state relative to the EZH2+/EGR1- state in PD 32 fibroblast cells. Hypergeometric P-value and expected overlap are indicated. **D)** Heatmap showing expression of overlapping mesenchymal drift genes identified in C) across all perturbations shown in Figure 1A.

**Figure S2: Expression of EGR1 and EZH2 target genes during neuronal maturation.** Heatmaps showing expression dynamics of all detected EGR1 and EZH2 target genes across neuronal maturation. This is a full-heatmap corresponding to Figure 2D.

**Figure S3. Expression dynamics during muscle stem cell activation.** A) Heatmap showing expression of representative marker genes across different states of muscle stem activation in young and aged donors. B-C) 2D kernel density plots of marker genes expression from single cells: MKI67: proliferation and MYOG for fate-specification and, MYOD1 as terminal differentiation marker.

FigureS1

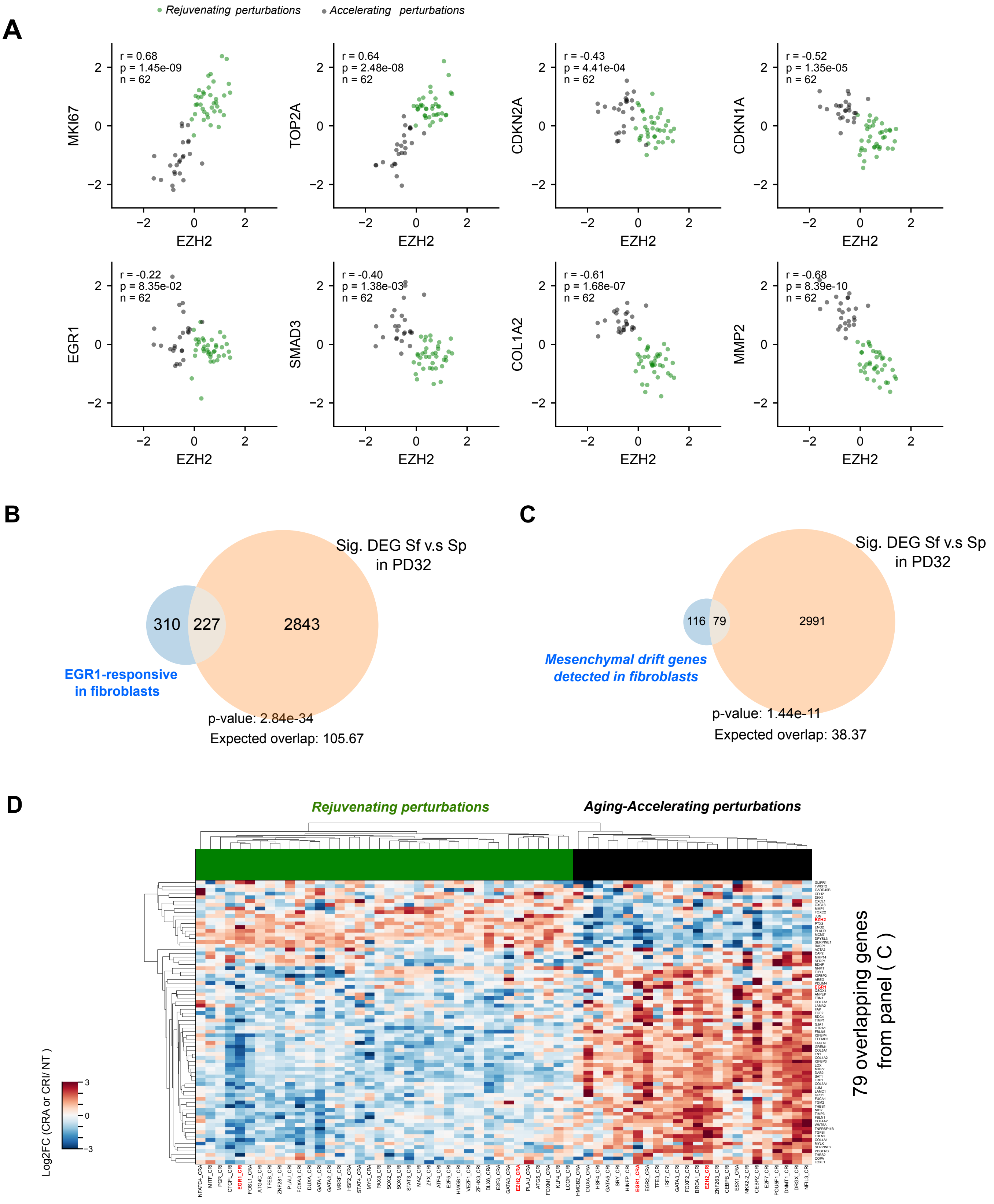
Rejuvenating perturbations
Accelerating perturbations
A
r = 0.68
p = 1.45e-09
n = 62
MKI67
EZH2
r = 0.64
p = 2.48e-08
n = 62
TOP2A
EZH2
r = -0.43
p = 4.41e-04
n = 62
CDKN2A
EZH2
r = -0.52
p = 1.35e-05
n = 62
CDKN1A
EZH2
r = -0.22
p = 8.35e-02
n = 62
EGR1
EZH2
r = -0.40
p = 1.38e-03
n = 62
SMAD3
EZH2
r = -0.61
p = 1.68e-07
n = 62
COL1A2
EZH2
r = -0.68
p = 8.39e-10
n = 62
MMP2
EZH2
B
Sig. DEG Sf v.s Sp
in PD32
310
227
2843
EGR1-responsive
in fibroblasts
p-value: 2.84e-34
Expected overlap: 105.67
C
Sig. DEG Sf v.s Sp
in PD32
116
79
2991
Mesenchymal drift genes
detected in fibroblasts
p-value: 1.44e-11
Expected overlap: 38.37
D
Rejuvenating perturbations
Aging-Accelerating perturbations
Log2FC (CRA or CRI/ NT )
79 overlapping genes
from panel ( C )

Figure S2

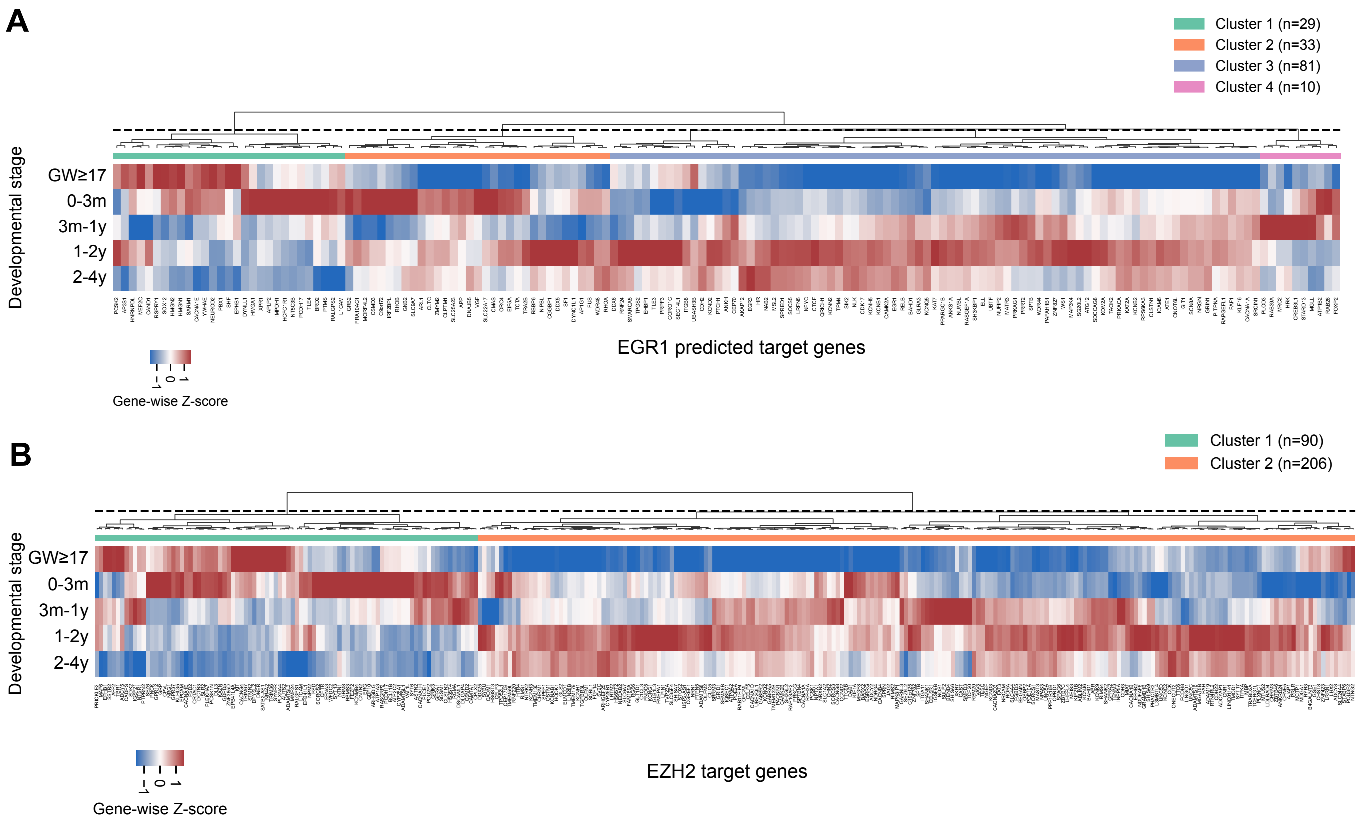
A
Cluster 1 (n=29)
Cluster 2 (n=33)
Cluster 3 (n=81)
Cluster 4 (n=10)
Developmental stage
GW≥17
0-3m
3m-1y
1-2y
2-4y
EGR1 predicted target genes
-1 0 1
Gene-wise Z-score
B
Cluster 1 (n=90)
Cluster 2 (n=206)
Developmental stage
GW≥17
0-3m
3m-1y
1-2y
2-4y
EZH2 target genes
-1 0 1
Gene-wise Z-score

### Figure S3

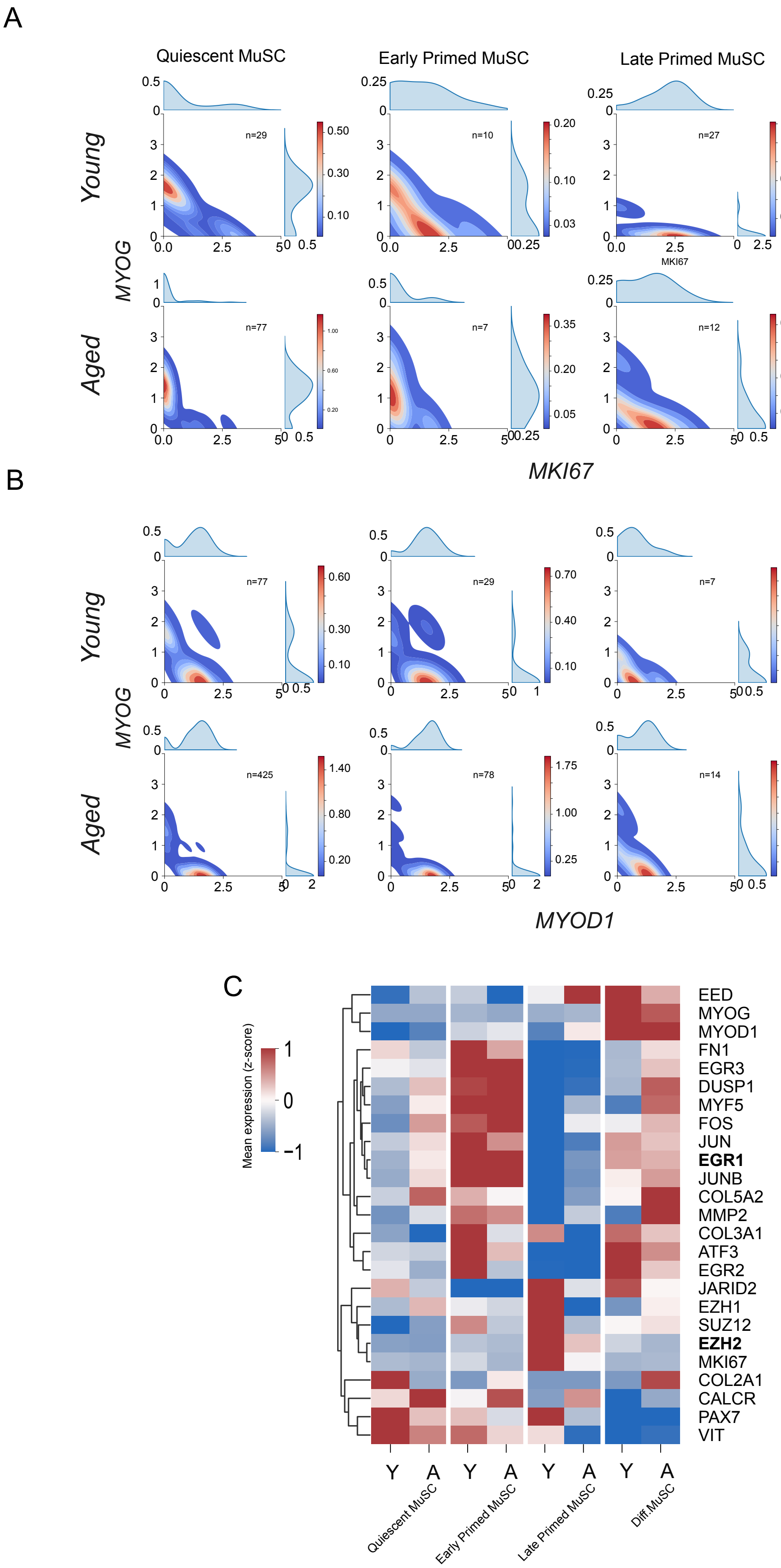
A
Quiescent MuSC
Early Primed MuSC
Late Primed MuSC
Diff.MuSC
Young
Aged
MYOG
MKI67
n=29
n=10
n=27
n=69
n=77
n=7
n=12
n=70
B
Young
Aged
MYOG
MYOD1
n=77
n=29
n=7
n=82
n=425
n=78
n=14
n=171
C
Mean expression (z-score)
EED
MYOG
MYOD1
FN1
EGR3
DUSP1
MYF5
FOS
JUN
EGR1
JUNB
COL5A2
MMP2
COL3A1
ATF3
EGR2
JARID2
EZH1
SUZ12
EZH2
MKI67
COL2A1
CALCR
PAX7
VIT
Y A Y A Y A Y A
Quiescent MuSC
Early Primed MuSC
Late Primed MuSC
Diff.MuSC
